# Coordinated control of the ADP-heptose/ALPK1 signalling network by the E3 ligases TRAF6, TRAF2/c-IAP1 and LUBAC

**DOI:** 10.1101/2022.07.19.500647

**Authors:** Tom Snelling, Natalia Shpiro, Robert Gourlay, Frederic Lamoliatte, Philip Cohen

## Abstract

ADP-heptose activates the protein kinase ALPK1 triggering TIFA phosphorylation at Thr9, the recruitment of TRAF6 and the subsequent production of inflammatory mediators. Here, we demonstrate that ADP-heptose also stimulates the formation of Lys63- and Met1-linked ubiquitin chains to activate the TAK1 and canonical IKK complexes, respectively. We further show that the E3 ligases TRAF6 and c-IAP1 operate redundantly to generate the Lys63-linked ubiquitin chains required for pathway activation, which we demonstrate are attached to TRAF6, TRAF2 and c-IAP1, and that c-IAP1 is recruited to TIFA by TRAF2. ADP-heptose also induces activation of the kinase TBK1 by a TAK1-independent mechanism, which requires TRAF2 and TRAF6. We establish that ALPK1 phosphorylates TIFA directly at Thr177 as well as Thr9 in vitro. Thr177 is located within the TRAF6-binding motif and its mutation to Asp prevents TRAF6 but not TRAF2 binding, indicating a role in restricting ADP-heptose signalling. We conclude that ADP-heptose signalling is controlled by the combined actions of TRAF2/c-IAP1 and TRAF6.

## Introduction

The early detection of microbial pathogens is mediated via germline-encoded pattern recognition receptors (PRRs), which detect abundant and evolutionarily conserved molecules present in microbial pathogens that are absent or inaccessible in host cells, called pathogen-associated molecular patterns (PAMPs) (Medzhitov, 2007). The activation of PRRs triggers intracellular signal transduction pathways, which culminate in the generation of cytokines that mediate the host response to infection.

ADP-L,D-heptose and ADP-D,D-heptose are metabolites required for the incorporation of L,D-heptose and D,D-heptose into the inner and outer core of bacterial lipopolysaccharide (LPS), respectively (Stein *et al*., 2017). D,D-heptose is also a constituent of S-layer glycoproteins present on the cell surface of archaea and bacteria (Schäffer and Messner, 2004), while septacidin and hygromycin B, which are secondary metabolites from Gram positive bacteria, require ADP-LD-heptose and ADP-DD-heptose, respectively, for their biosynthesis (Tang *et al*., 2018). In 2018, ADP-L,D-heptose and ADP-D,D-heptose (hereafter, ADP-heptose) were identified as activators of the atypical mammalian alpha-protein kinase 1 (ALPK1). The binding of ADP-heptoses to an N-terminal domain of ALPK1 induces an allosteric transition that activates the C-terminal kinase domain (Zhou *et al*., 2018). These investigators also showed that ADP-heptoses induced the activation of the transcription factor NF-κB and *IL-8* gene transcription in human embryonic kidney (HEK) 293 cells by a pathway that requires the expression of TNF receptor-activating factor (TRAF)-interacting protein with forkhead-associated (FHA) domain (TIFA) and TRAF6.

TIFA (comprising 184 amino acid residues) is a human FHA-containing protein that was identified in 2002 as a TRAF2-binding protein (T2BP) during a mammalian two-hybrid screen. Its overexpression in HEK293 cells was shown to activate the transcription factors NF-κB and AP-1 (Kanamori *et al*., 2002), and it was proposed to be a component of the TNF signalling pathway in which TRAF2 has a key role alongside TRAF5 (Tada *et al*., 2001). Subsequently, T2BP was shown to interact with TRAF6 and renamed TIFA (Takatsuna *et al*., 2003). Based on overexpression studies, TIFA was suggested to function as a scaffolding protein operating between IRAK1 and TRAF6 in MyD88-dependent signalling pathways (Takatsuna et al., 2003) that are triggered by Interleukin-1 (IL-1) family members and by ligands that activate Toll-Like Receptors (TLRs). TIFA was also shown to stimulate the oligomerisation of TRAF6 and the activation of its E3 ligase activity (Ea *et al*., 2004).

FHA domains bind to phospho-threonine residues in proteins (Mahajan *et al*., 2008). To identify sites of phosphorylation in TIFA, Huang and co-workers overexpressed FLAG-TIFA in HEK293 cells and after digestion with a cocktail of proteases that included GluC, as described elsewhere (Fang *et al*., 2010), identified Thr9 as one of several amino acid residues in TIFA that were phosphorylated (Huang *et al*., 2012).Wild type (WT) TIFA was shown to undergo self-oligomerisation through an intermolecular interaction between the FHA domain of one TIFA molecule and phospho-Thr9 of another; this did not occur if WT TIFA was replaced by the TIFA[T9A] mutant (Huang *et al*., 2012). Other investigators also identified Thr9 as an amino acid residue in TIFA that became phosphorylated in response to heptose-1,7-bisphosphate (a precursor of ADP-heptose) in Jurkat cells (Gaudet *et al*., 2015). At this time, the identity of the protein kinase that phosphorylated TIFA was unknown. Later, the ADP-heptose-dependent activation of NF-κB and *IL-8* gene transcription was found to be abolished when WT TIFA was replaced by the TIFA[T9A] mutant (Zhou *et al*., 2018).

A variety of gram negative bacteria, including *Shigella flexneri*, *Salmonella enterica S*erovar Typhimurium, *Neisseria meningitidis*, and *Helicobacteria pylori* induce the oligomerisation of TIFA in the gastric epithelial cell line AGS (Gall *et al*., 2017; Milivojevic *et al*., 2017; Stein *et al*., 2017; Zimmermann *et al*., 2017). The cytokine deficiency-induced colitis susceptibility locus (*Cdcs1*), which is linked to mouse inflammatory bowel disease, also contains the *alpk1* gene. The *alpk1* gene is present in the genetic interval termed ‘Hiccs’ on mouse chromosome 3 that regulates *Helicobacter hepaticus*-induced colitis in a RAG1 KO background and leads to the eventual onset of colitis-associated colorectal cancer (Boulard *et al*., 2012). RAG1 KO mice develop colitis spontaneously upon infection with *H. hepaticus*, which is exacerbated in RAG1/ALPK1 double KO mice (Ryzhakov *et al*., 2018). Intriguingly, the *alpk1* and *tifa* genes are adjacent on mouse chromosome 3, and the *ALPK1* and *TIFA* genes are adjacent on human chromosome 4. Here we have performed a detailed biochemical and genetic analysis of the events that take place during the first hour of ADP-heptose-ALPK1-TIFA-signalling, which has revealed that TRAF2/c-IAP1 and TRAF6 both have critical roles in the regulation of this network.

## Results

### The ADP-heptose stimulated activation of MAP kinases and the canonical IKK complex requires ALPK1 and TIFA

In previous studies, the ADP-LD-heptose signalling pathway has been monitored in human embryonic kidney 293 (HEK293) cells by the activation of the transcription factor NF-κB using a luciferase reporter assay and by measuring *IL-8* transcript levels 4 h after the electroporation of 10 µM ADP-LD-heptose or the extracellular addition of 100 µM ADP-LD-heptose (Zhou *et al*., 2018). Here we monitored pathway activation routinely by the phosphorylation (activation) of the mitogen-activated protein kinases (MAPKs) p38α MAPK, p38γ MAPK and c-Jun N-terminal kinases 1 and 2 (JNK1/JNK2), and by the phosphorylation (activation) of the canonical IκB kinase (IKK) complex and its substrate NF-κB1 (Beinke *et al*., 2004; Waterfield *et al*., 2004), and by the disappearance of its substrate IκBα. The IKKβ-catalysed phosphorylation of IκBα triggers its ubiquitylation by the E3 ligase SCF-β^TRCP^ causing its proteasomal degradation and the activation of NF-κB (Yamamoto *et al*., 2000). We found that ADP-DD-heptose or ADP-LD-heptose activated the pathway rapidly and near maximally when included in the culture medium at 10 μM (Fig S1). ADP-DD-heptose (hereafter called ADP-heptose) was used for all subsequent experiments.

Pathway activation in HEK293 cells or primary mouse bone marrow-derived macrophages (BMDM) could be detected within 10 min and was maximal after 20-30 min. The strength of pathway activation induced by ADP-heptose was similar to that induced by IL-1β in HEK293 cells (Fig 1A) or by R848, an activator of Toll-Like Receptor 7 (TLR7), in BMDM (Fig 1B). The rapid ADP-heptose-stimulated destruction of IκBα was followed by its resynthesis within 45-60 min, because IκBα is an NF-κB-dependent gene (Fig 1A).

**Figure 1.**
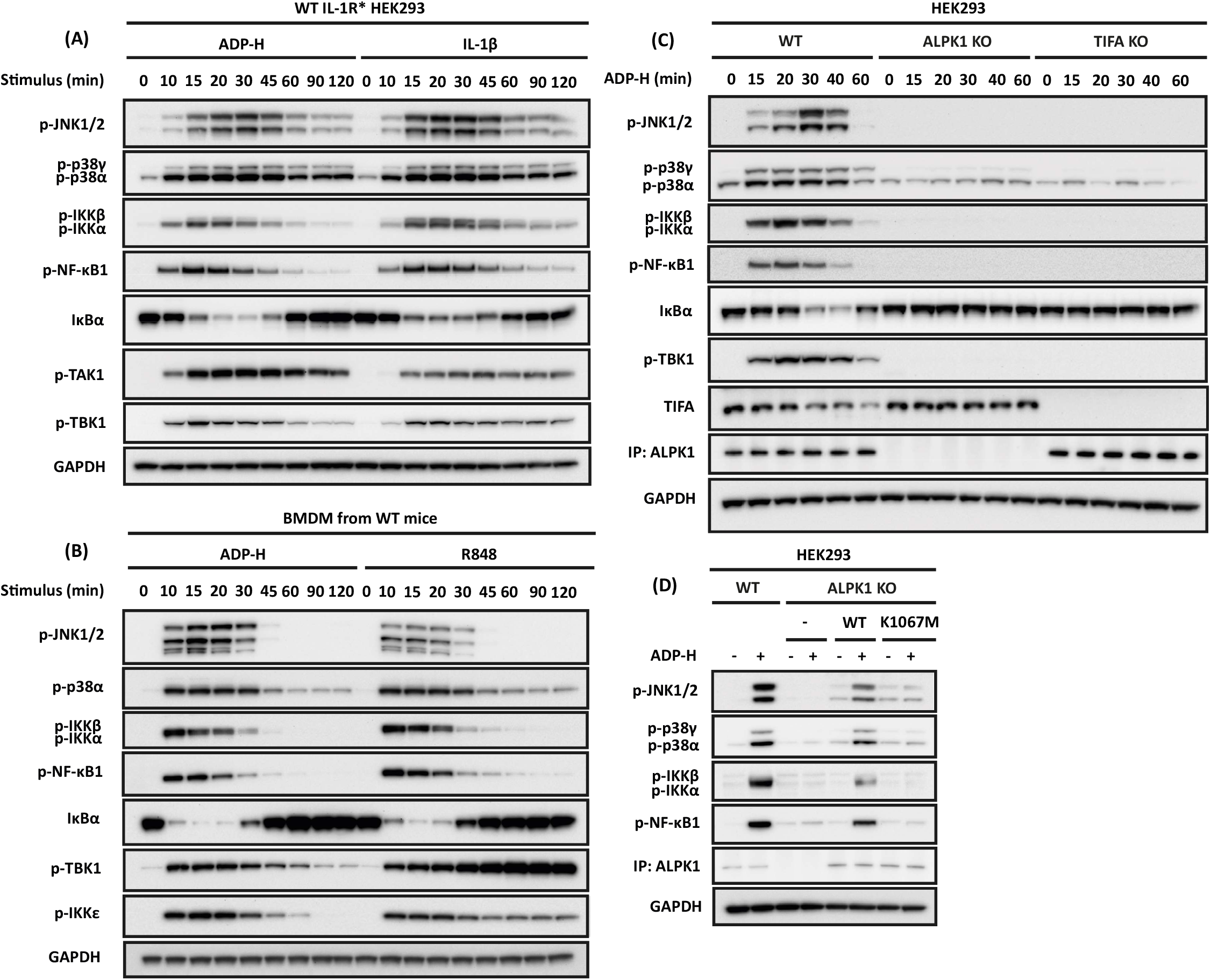
ADP-heptose induces the ALPK1- and TIFA-dependent activation of MAP kinases and the canonical IKK complex. **(A, B)** IL-1R* HEK293 cells (**A**) or BMDM from wild type (WT) mice (**B**) were stimulated for the times indicated with ADP-heptose (ADP-H) or IL-1β (**A**) and ADP-H or R848 **(B)**. **(C)** Parental, ALPK1 KO and TIFA KO HEK293 cells were stimulated for the times indicated with ADP-H **(D)** ALPK1 KO HEK293 cells were transfected with empty vector (-), FLAG-ALPK1 (WT) or FLAG-ALPK1[K1067M] (K1067M) and 48 h later were stimulated for 20 min with ADP-H. **(A-D)** Cell extracts were analysed directly by SDS-PAGE and immunoblotting was performed using antibodies that recognise phosphorylated (p) forms of the indicated proteins, or with antibodies recognising all forms of TIFA, ALPK1 IκBα and GAPDH. In C and D, ALPK1 was detected after its immunoprecipitation from the extracts with an ALPK1 antibody (see methods).

As expected, the knockout (KO) of ALPK1 or TIFA in HEK293 cells abolished ADP-heptose signalling (Fig 1C) but had no effect on IL-1β signalling (Fig S2A). It has been suggested that TIFA acts as a scaffold between IRAK1 and TRAF (Takatsuna *et al*., 2003), but the KO of TIFA had no effect or IL-1β signalling (Fig 1C) and the KO of IRAK1 had no effect on ADP-heptose signalling (Fig S2B). ADP-heptose signalling was restored by re-transfecting wild type (WT) ALPK1 into ALPK1 KO cells, but not by re-transfection of the kinase-inactive ALPK1[K1067M] mutant (Fig 1D).

### The expression of TAK1 and its protein kinase activity is required for ADP-heptose signalling in HEK293 cells

In HEK293 cells, IL-1β signalling requires the protein kinase TAK1 (also termed MAP3K7). TAK1 activates the MAPK kinases (MKKs) that activate p38α/γ and JNK1/2 and also initiates IKK activation (Zhang *et al*., 2014; Strickson *et al*., 2017), but the MAP3K that mediates ADP-heptose signalling is unknown. We found that ADP-heptose induced the phosphorylation of TAK1 at Thr187 in the activation loop (Fig 1A) and therefore investigated whether TAK1 was the MAP3K that drives this pathway. ADP-heptose signalling was abolished in TAK1 KO HEK293 cells (Fig 2A) and could be restored by the re-expression of WT TAK1 but not by the kinase-inactive TAK1[D175A] mutant (Fig 2B). ADP-heptose signalling was also abolished by two relatively specific, structurally unrelated inhibitors of TAK1, 5(Z)-7-oxozeanol (Ninomiya-Tsuji *et al*., 2003) (Fig 2C) and NG25 (Tan *et al*., 2015) (Fig 2D) whose off-target effects on other protein kinases do not overlap (https://www.kinase-screen.mrc.ac.uk/kinase-inhibitors). We also found that both TAK1 inhibitors suppressed the phosphorylation of TAK1 at Thr187 (Fig 2C and 2D), implying that TAK1 is being activated by autophosphorylation.

**Figure 2.**
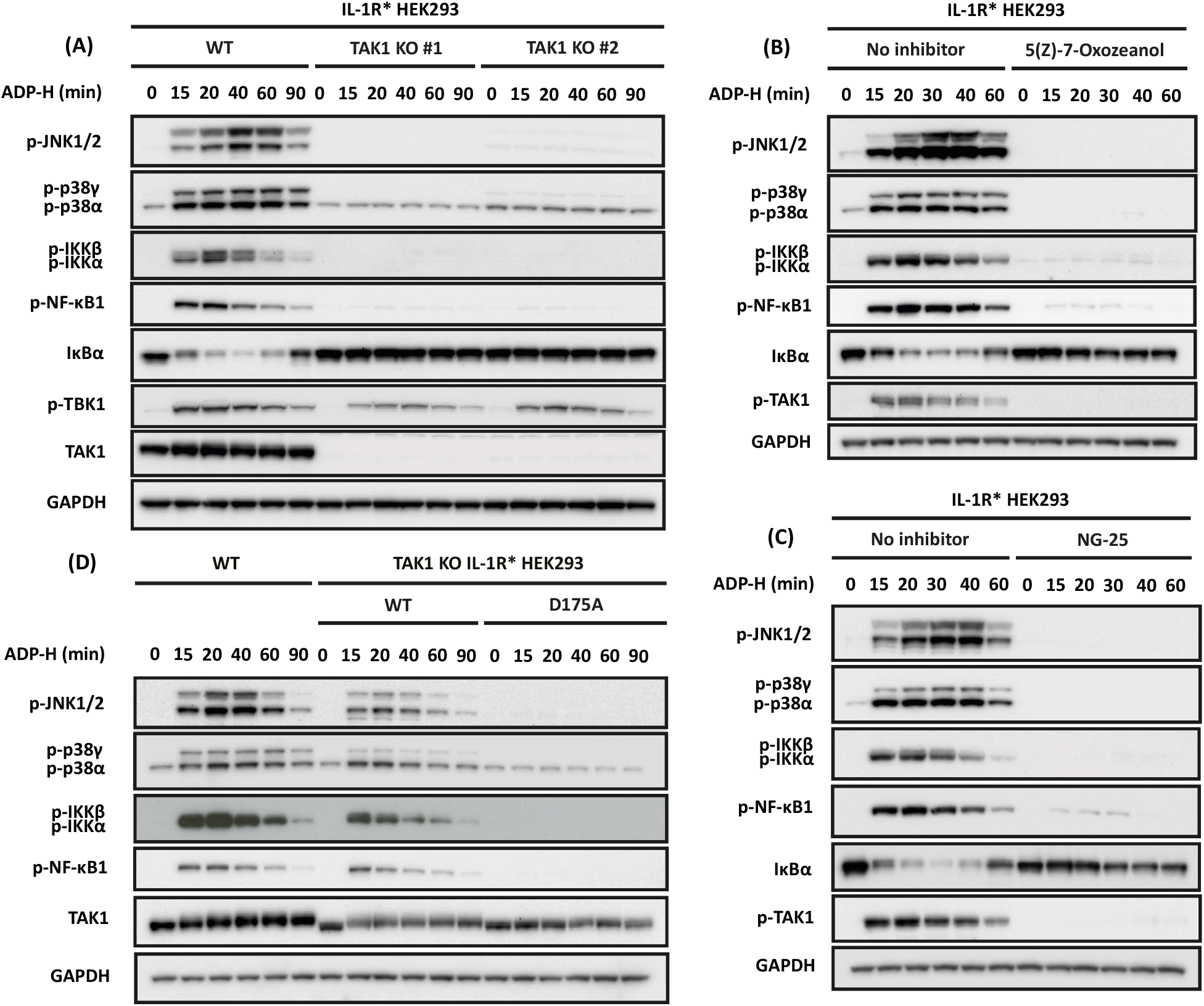
TAK1 and its protein kinase activity are required for ADP-heptose to activate MAP kinases and the canonical IKK complex. **(A)** The parental and two independently isolated clones of TAK1 KO IL-1R* HEK293 cells were stimulated for the times indicated with ADP-heptose (ADP-H). **(B)** As in (A), except that WT TAK1 or the kinase-inactive TAK1[D175A] mutant were stably expressed in TAK1 KO cells under a doxycycline-inducible promoter and expression induced for 16 h with 50 ng/ml doxycycline, prior to stimulation with ADP-H for the times indicated. **(C, D)** IL-1R* HEK293 cells were treated for 1 h with 2 µM (5Z)-7-oxozeaenol (**C**) or 5 µM NG25 (**D**) and then stimulated with ADP-H for the times indicated. **(A-C)** Cell extracts were subjected to SDS-PAGE and immunoblotted with the antibodies indicated.

### ADP-heptose stimulates the formation of Lys63-linked and Met1-linked ubiquitin chains

In the IL-1β signalling pathway TAK1 is activated by the formation of Lys63-linked ubiquitin (K63-Ub) chains, which bind to the TAB2 and TAB3 regulatory components of the TAK1 complex, inducing a conformational change that permits the autoactivation of TAK1 (Kanayama *et al*., 2004; Kulathu *et al*., 2009; Zhang *et al*., 2014). IL-1β signalling also stimulates the formation of Met1-linked ubiquitin (M1-Ub) chains in HEK293 cells, which are catalysed by the HOIP component of the Linear Ubiquitin Assembly Complex (LUBAC) (Kirisako *et al*., 2006; Tokunaga *et al*., 2009). The M1-Ub chains bind to the NEMO (NF-κB essential modifier) subunit of the canonical IKK complex (Lo *et al*., 2009; Rahighi *et al*., 2009), inducing a conformational change that enables TAK1 to initiate the activation of IKKβ (Zhang *et al*., 2014). Here, we found that ADP-heptose also stimulates the formation of K63-Ub and M1-Ub chains in HEK293 cells (Fig S3A) or mouse BMDM (Fig S3B), although the extent of ubiquitylation is lower than in IL-1β-stimulated HEK293 cells (Fig S3A) or R848-stimulated mouse BMDM (Fig S3B). Consistent with a key role for K63-Ub chains in the activation of TAK1, we found that ADP-heptose signalling was suppressed in TAB2/3 double KO HEK293 cells, even allowing for the decreased expression of TAK1 in these cells (Fig S3C). In addition, phosphorylation of NF-κB1 was reduced in mouse embryonic fibroblasts (MEFs) from mice expressing the catalytically inactive HOIP mutant, HOIP[C879S] (Fig S3D) and obliterated in MEFs from mice expressing a ubiquitin-binding defective mutant of NEMO, NEMO[D311N] (Fig S3E), without affecting the phosphorylation of p38α (Fig S3D and S3E).

### TRAF6 is required for ADP-heptose signalling, but its E3 ligase activity is not

The finding that the expression of TRAF6 is required for ADP-heptose signalling (Zhou *et al*., 2018) was confirmed in the present study, by showing that MAPK and IKK activation were abolished in TRAF6 KO HEK293 cells (Fig 3A) or in primary TRAF6 KO macrophages derived from foetal liver cells (Fig 3B). Interestingly, ADP-heptose signalling could be restored to TRAF6 KO HEK293 cells not only by the re-expression of WT TRAF6, but also by the re-expression of the E3 ligase-inactive TRAF6[L74H] mutant or the TRAF6[120-522] mutant in which the RING (Really Interesting New Gene) domain of TRAF6, which carries the E3 ligase activity, is deleted (Fig 3C). The ADP-heptose-stimulated activation of MAPKs was similar in primary BMDM from knock-in mice expressing the TRAF6[L74H] mutant and in WT mice (Fig 3D). Although the phosphorylation of IKKα/β was slightly reduced in BMDM from TRAF6[L74H] mice, the rate of degradation of IκBα and its rate of resynthesis were similar to BMDM from WT mice (Fig 3D), indicating that the IKK activity was still sufficient to activate NF-κB near maximally. Taken together these experiments established that although TRAF6 is required for ADP-heptose signalling, its E3 ligase activity is not.

**Figure 3.**
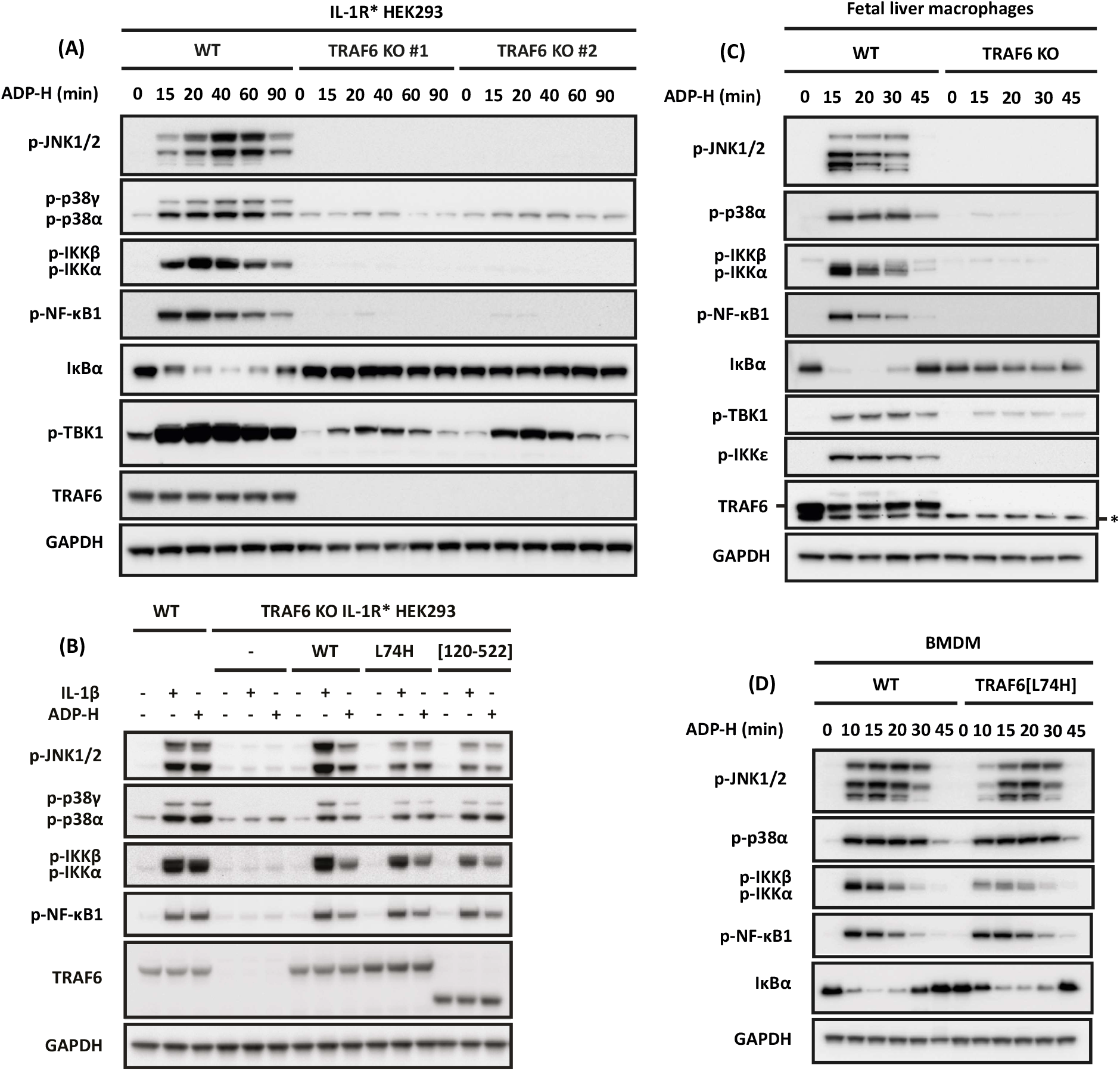
TRAF6 is required for ADP-heptose signalling in HEK293 cells and primary macrophages, but its E3 ligase activity is not. **(A)** Parental and two independently isolated clones of TRAF6 KO IL-1R* HEK293 cells were stimulated for the times indicated with ADP-heptose (ADP-H). **(B)** Foetal liver macrophages from wild type (WT) and TRAF6 KO embryos were stimulated for the times indicated with ADP-H. **(C)**, As in (A), except that WT TRAF6, TRAF6[L74H] or TRAF6[120-522] were stably expressed in TRAF6 KO IL-1R* HEK293 cells under a constitutive promoter and stimulated for 20 min with ADP-H or IL-1β. **(D)** BMDM from WT and TRAF6[L74H] mice were stimulated for the times indicated with ADP-H. **(A-D)** Cell extracts were subjected SDS-PAGE and immunoblotted using the indicated antibodies. An asterisk indicates protein(s) recognised non-specifically by an antibody.

The TRAF6 E3 ligase activity is also not essential for IL-1β signalling in HEK293 cells or TLR signalling in primary BMDM (Strickson *et al*., 2017). This is because the Pellino E3 ligases (Pellino 1 and Pellino 2), which are activated by an IRAK1-catalysed phosphorylation mechanism during IL-1β signalling, also form K63-Ub chains (Ordureau *et al*., 2008; Smith *et al*., 2009; Goh *et al*., 2012) and operate redundantly with TRAF6 to generate the K63-Ub chains required for the activation of TAK1 (Cohen and Strickson, 2017; Strickson *et al*., 2017). We therefore wondered whether the K63-Ub chains required for the ADP-heptose-stimulated activation of TAK1 might also be formed by TRAF6 and another E3 ligase, explaining why the E3 ligase activity of TRAF6 was not essential for ADP-heptose signalling.

### Identification of proteins that immunoprecipitate with TRAF6 in ADP-heptose-stimulated cells

To investigate the identity of the putative E3 ligase, we studied the composition of TRAF6 complexes formed after stimulation for 20 min with ADP-heptose, the time at which IKK activation was maximal. We stably expressed FLAG-TRAF6 in TRAF6 KO HEK293 cells, captured TRAF6 by immunoprecipitation with a FLAG antibody, and released the TRAF6-binding proteins by incubation with FLAG peptide. These proteins were then identified by mass spectrometry (MS) using Tandem Mass Tag (TMT) quantification. The Volcano plot (Fig 4A) revealed that major proteins captured in an ADP-heptose-dependent manner included TIFA, the four components of the TAK1 complex (TAB1, TAB2, TAB3 and TAK1), IKKα and IKKβ, the three components of LUBAC (Sharpin, HOIL-1 and HOIP) and the HOIP-interacting protein WRNIP1 that contains a ubiquitin-binding zinc finger domain and a Glu-Glu-His-Tyr-Asn motif that may mediate binding to the PUB domain of HOIP (Schaeffer *et al*., 2014). They also included A20 and ABIN1 (A20-binding inhibitor of NF-κB), two proteins known to restrict IL-1β and TLR signalling by binding to M1-Ub and K63-Ub chains (Cohen and Strickson, 2017). Other proteins that were captured included TRAF2, its known binding partner c-IAP1, and the c-IAP1-binding proteins SMAC and HtrA2/Omi. SMAC and HtrA2 are mitochondrial proteins that bind to c-IAP1 only when mitochondria are disrupted (Duckett, 2005), which most likely occurred in the present study during cell lysis, as a result of the inclusion of the detergent Triton X-100 in the cell lysis buffer.

**Figure 4.**
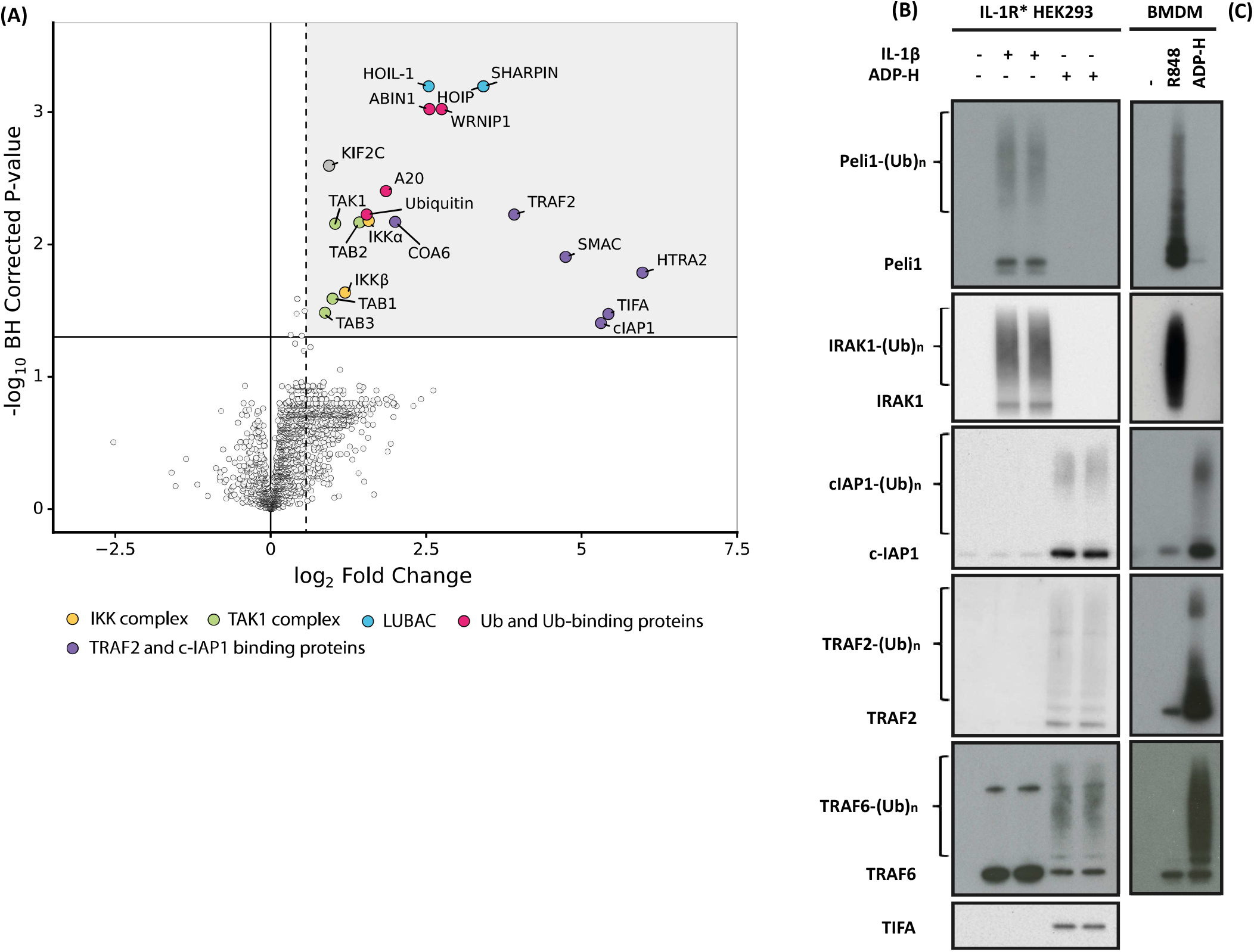
Proteins enriched in TRAF6 immunoprecipitates in ADP-heptose-stimulated HEK293 cells. **(A)** IL-1R* HEK293 cells were stimulated for 20 min with ADP-heptose (ADP-H) and proteins in cell extracts immunoprecipitated with FLAG-TRAF6 were analysed by TMT mass spectrometry. The figure shows a volcano plot identifying the proteins where enrichment attained statistical significance and where the fold-increase was >1.5-fold. **(B)** HEK293 cells were stimulated for 15 min with ADP-H or IL-1β and the proteins shown that were captured from the cell extracts on Halo-NEMO beads were identified by SDS-PAGE followed by immunoblotting with the antibodies indicated. **(C)** WT BMDM were stimulated for 15 min with ADP-H or R848 and analysed with the same antibodies used in (**B**), except for cIAP1, where a mouse-specific antibody was used (see methods). Ubiquitylated forms of proteins are indicated by (Ub)_n_.

TRAF2 was originally found to bind to TIFA in a mammalian two-hybrid screen (Kanamori *et al*., 2002) and in AGS cells infected with *H. pylori* (Zimmermann *et al*., 2017), and c-IAP1 is an E3 ligase. We therefore wondered whether TRAF2 might recruit c-IAP1 to TIFA during ADP-heptose signalling to produce K63-Ub chains required for the activation of TAK1. To investigate this hypothesis, we used Halo-NEMO beads to capture K63- and M1-Ub chains from HEK293 cell extracts, as well as the proteins to which they are attached covalently and non-covalently (Emmerich *et al*., 2013; Strickson *et al*., 2017). We found that Halo-NEMO beads captured ubiquitylated forms of c-IAP1, TRAF2 and TRAF6 in an ADP-heptose-dependent manner but did not capture IRAK1 or Pellino 1. In contrast, the Halo-NEMO beads captured ubiquitylated forms of Pellino 1 and IRAK1 in an IL-1β-dependent manner but did not capture c-IAP1 or TRAF2. The c-IAP2 isoform was not detected in the Halo-NEMO pulldowns because it is not expressed in HEK293 cells (Fig S4). The Halo-NEMO beads also captured ubiquitylated c-IAP1 and TRAF2 from the extracts of ADP-heptose stimulated mouse BMDM, but not Pellino 1 or IRAK1; conversely, Halo-NEMO beads captured ubiquitylated Pellino 1 and IRAK1 from the extracts of R848-stimulated mouse BMDM but not c-IAP1 or TRAF2 (Fig 4C). TRAF6 was also captured from the extracts of IL-1β-stimulated HEK293 cells (Fig 4B) and R848-stimulated BMDM (Fig 4C) but, in contrast to ADP-heptose-stimulated cells, it was not ubiquitylated and was presumably captured by Halo-NEMO beads because of its interaction with ubiquitylated IRAK1. TIFA was also captured by Halo-NEMO beads from the extracts of ADP-heptose-stimulated HEK293 cells but was not ubiquitylated (Fig 4B).

Taken together, these observations indicated that ADP-heptose did not induce the activation and autoubiquitylation of Pellino 1, suggesting that Pellino isoforms were unlikely to be the putative E3 ligases operating redundantly with TRAF6 during ADP-heptose signalling.

### The TRAF6 and c-IAP1 E3 ligases generate the K63-Ub chains needed to propagate the ADP-heptose signal

To investigate the role of TRAF2/c-IAP1 in ADP-heptose signalling, we stably re-expressed WT TRAF6 or TRAF6[L74H] in TRAF2/TRAF6 double KO HEK293 cells. The ADP heptose-stimulated phosphorylation of IKKα/β, NF-κB1, JNK1/2 and p38α/γ MAPK was restored to TRAF6 KO cells by the re-expression of either WT TRAF6 or TRAF6[L74H], as expected, but IKKα/β, NF-κB1 and p38γ MAPK activation could only be restored by WT TRAF6 in TRAF2/TRAF6 double KO cells (Fig 5A). However, trace phosphorylation of JNK1/2 and p38α MAPK activation was restored to TRAF2/TRAF6 double KO cells by the re-expression of TRAF6[L74H]. We speculate that this may be driven by trace amounts of TRAF5 present in these cells.

**Figure 5.**
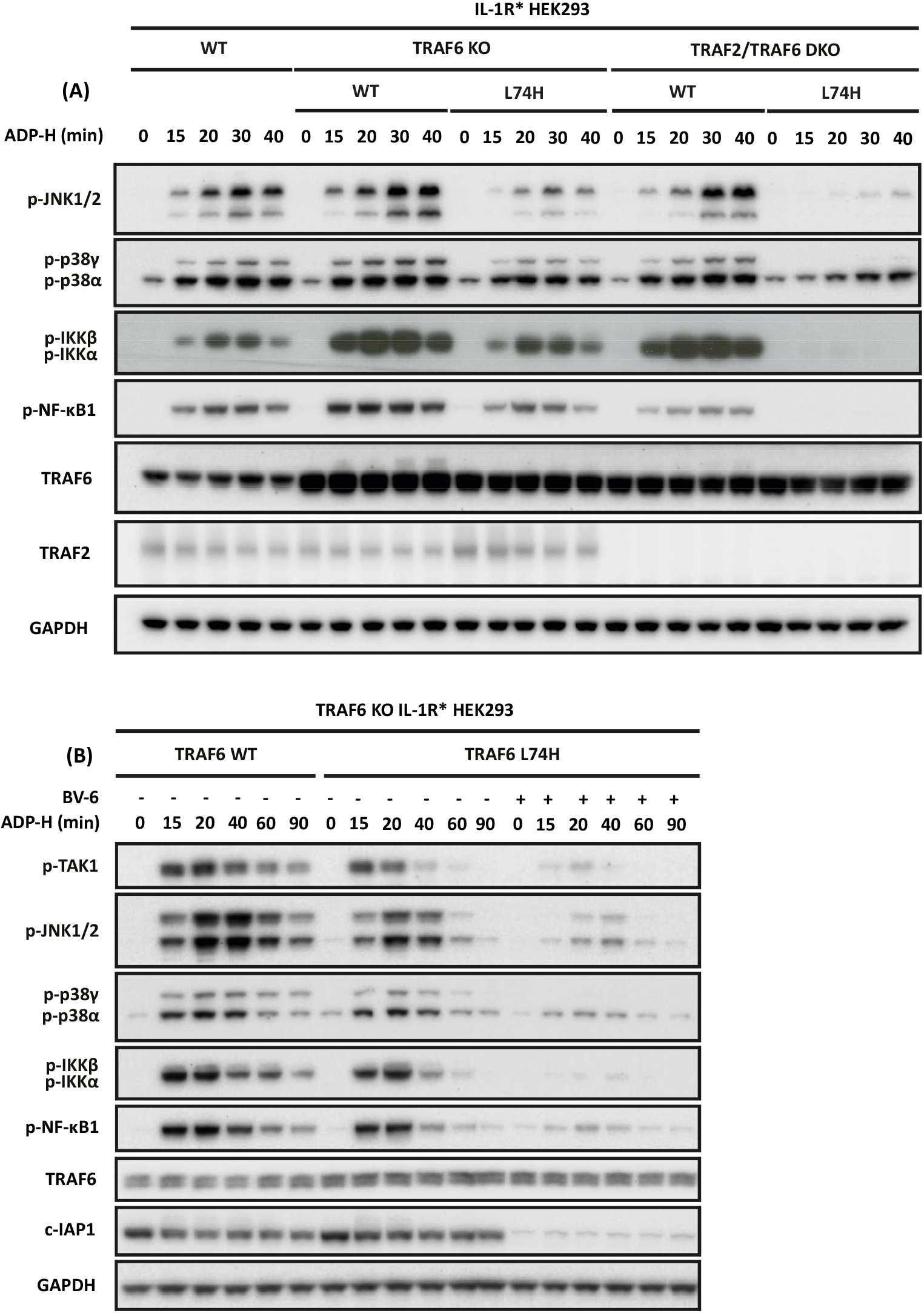
The TRAF6 and TRAF2/c-IAP1 operate redundantly to generate the ubiquitin chains required for ADP-heptose signalling. **(A)** Parental, TRAF6 KO or TRAF2/6 double KO IL-1R* HEK293 cells in which WT TRAF6 or TRAF6[L74H] had been stably re-expressed under a constitutive promoter were stimulated for the times indicated with ADP-H. **(B)** TRAF6 KO IL-1R* HEK293 cells in which WT TRAF6 or TRAF6[L74H] had been stably re-expressed as in (**A**) were treated for 5 h with 5 µM BV-6 prior to stimulation for the indicated times with ADP-H. **(A, B)** The cell extracts were subjected to SDS-PAGE and immunoblotting was performed with the antibodies indicated.

Since TRAF2 appears to be an inactive “pseudo” E3 ligase (Yin *et al*., 2009), these results were consistent with the notion that a key role of TRAF2 is to recruit c-IAP1 into the TIFA signalling complex, where it acts redundantly with TRAF6 to produce the K63-Ub chains required to activate TAK1. We tested this hypothesis by studying the effect of the SMAC mimetic BV-6 (Li *et al*., 2011). SMAC mimetics are small chemical entities that bind to IAP family members, inducing their autoubiquitylation and proteasomal destruction (Wang, 2010). Incubation of HEK293 cells with BV-6 greatly reduced the expression of c-IAP1, as expected, and greatly reduced ADP-heptose signalling in TRAF6 KO cells reconstituted with TRAF6[L74H] (Fig 5B). The residual cIAP1 expression may explain why signalling was not completely abolished.

Taken together, these results support the conclusion that c-IAP1 is the E3 ligase that operates redundantly with TRAF6 to generate the K63-Ub chains required to activate TAK1 and hence the activation of JNK1/2 and IKKα/β and the phosphorylation of the IKKβ substrate NF-κB1 during ADP-heptose signalling in HEK293 cells.

### The effects of TRAF2/c-IAP1 are not mediated by the non-canonical NF-κB pathway

It is well established that TRAF2-cIAP1/2 complexes are components of the non-canonical NF-κB pathway that induces the expression of NF-κB-inducing kinase (NIK) (Sun, 2017). It has also been reported that ADP-heptose activates the non-canonical NF-κB pathway in AGS cells (Maubach *et al*., 2021), and we confirmed this result in HT-1080 cells (Fig S5B). However, the ADP-heptose induced expression of NIK was only detected after 2 h, much later than the degradation of IκBα, which was complete by 20 min. Moreover, even in HT-1080 cells, the ADP-heptose-stimulated appearance of NIK was much weaker than that induced by the TNF superfamily member TWEAK (TNF-like weak inducer of apoptosis) (Fig S5B). More importantly, we were unable to detect any appearance of NIK in HEK293 cells after stimulation with ADP-heptose for up to 8 h (Fig S5A), indicating that ADP-heptose does not activate the non-canonical NF-κB pathway in HEK293 cells. The rapid effects of TRAF2/c-IAP1 on ADP-heptose signalling in HEK293 cells described in the present study are therefore independent of the non-canonical NF-κB signalling pathway.

### The ADP heptose-stimulated activation of TBK1 is mediated by TRAF6 and TRAF2

The protein kinase TANK-binding kinae-1 (TBK1) is an IKK-related kinase that is converted from an inactive to an active form by phosphorylation at Ser172 (Kishore *et al*., 2002). In the present study we found that the phosphorylation (activation) of TBK1 was induced by ADP-heptose in HEK293 cells (Fig 1A) and BMDM (Fig 1B) but not in ALPK1 or TIFA KO HEK293 cells (Fig 1C). The ADP-heptose-stimulated phosphorylation of TBK1 was unaffected in TAK1 KO cells (Fig 2A), reduced in TRAF6 KO cells (Fig 3A) and abolished in TRAF2/TRAF6 double KO HEK293 cells (Fig S5A). We did not study the regulation of IKKε, the other IKK-related kinase, because its expression was extremely low in HEK293 cells (Fig S4). However, we did detect the phosphorylation of IKKε in BMDM (Fig 1B), which was disrupted in TRAF6 KO foetal liver macrophages (Fig 3C). The role of TBK1 in this pathway is considered further in the discussion.

### ALPK1 phosphorylates TIFA at Thr9 and Thr177

The mutation of Thr9 to Ala was shown to abolish TIFA oligomerisation in overexpression studies before the ADP-heptose signalling pathway was discovered (Huang *et al*., 2012). Later, the replacement of TIFA by the TIFA[T9A] mutant was found to abolish ADP-heptose-stimulated NF-κB-dependent gene transcription and IL-8 transcript formation (Zhou *et al*., 2018). These experiments suggested that the phosphorylation of TIFA at Thr9 was required for ADP-heptose signalling but the direct phosphorylation of TIFA at Thr9 by ALPK1 *in vitro* has never been demonstrated. Therefore, the possibility that ALPK1 phosphorylates and activates another protein kinase, which then phosphorylates TIFA at Thr9, has also not been excluded.

We re-expressed WT and mutant forms of FLAG-TIFA in TIFA KO HEK293 cells. ADP-heptose induced the activation of JNK and IKK in cells re-expressing WT FLAG-TIFA, but not in cells expressing TIFA[T9A], as expected (Fig 6A). More surprisingly, ADP-heptose failed to induce signalling in cells expressing the TIFA[T9S] mutant, or in cells expressing the TIFA [T9D] or TIFA[T9E] mutants in which a negative charge was introduced to try to mimic the effect of phosphorylation (Fig 6A). Consistent with these findings, ADP-heptose failed to induce the binding of TRAF6 or TRAF2 to TIFA in cells expressing TIFA[T9A], TIFA[T9S], TIFA [T9D] or TIFA[T9E] (Fig 6B). We therefore studied the phosphorylation of TIFA by ALPK1 *in vitro* using purified proteins and Mg-[γ^32^P]ATP.

**Figure 6.**
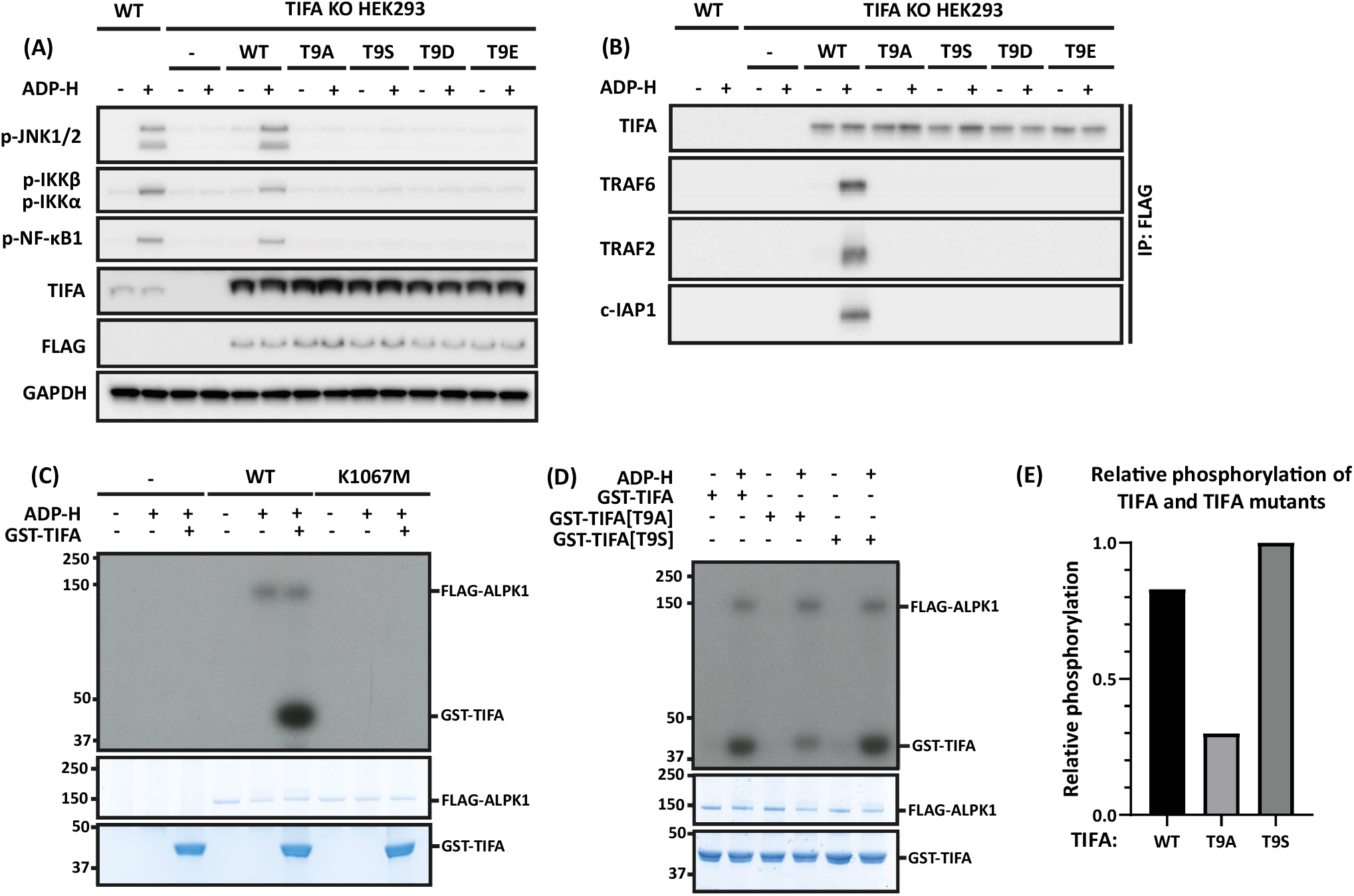
ALPK1 phosphorylates TIFA directly at Thr9 in vitro. **(A)** TIFA KO HEK293 cells were re-transfected with 500 ng of DNA plasmid encoding FLAG-TIFA or the indicated FLAG-TIFA mutants. 24 h later, the cells were stimulated for 20 min with ADP-heptose (ADP-H) and the cell extracts subjected to SDS-PAGE and immunoblotting with the antibodies indicated. **(B)** The cell extracts (1.0 mg protein) from (**A**) were immunoprecipitated with anti-FLAG and then subjected to SDS-PAGE and immunoblotting with the antibodies indicated **(C-E)** ALPK1 KO HEK293 cells were re-transfected with 2.5 μg plasmid DNA encoding empty vector (-), FLAG-ALPK1 and FLAG-ALPK1[K1067M] as indicated. 48 h later, ALPK1 was immunoprecipitated from the cell extracts with FLAG antibody and phosphorylation performed for 20 min in the presence or absence of 10 nM ADP-H and 8 µM GST-TIFA or GST-TIFA mutants and 0.1 mM [γ−^32^P]ATP (1000 cpm/pmol). Incorporation of ^32^P-radioactivity was visualised by autoradiography (**C, D**) and by Cerenkov counting (**E**).

FLAG-tagged WT ALPK1 phosphorylated itself and purified GST-TIFA *in vitro* but no incorporation of ^32^P-radioactivity into these proteins occurred when WT ALPK1 was replaced by the kinase-inactive ALPK1[K1067M] mutant (Fig 6C). The phosphorylation of TIFA was reduced by 64% when GST-TIFA was replaced by GST-TIFA[T9A] (Fig 6D and 6E), suggesting that Thr9 was not the only site of phosphorylation. Interestingly, ALPK1 phosphorylated GST-TIFA[T9S] as efficiently as WT TIFA (Fig 6D and 6E), indicating that TIFA phosphorylated at Ser9 is unable to interact with the FHA domain of TIFA and hence cannot induce the oligomerisation of TIFA that is required to initiate signalling.

Inspection of the amino acid sequence of human TIFA revealed that the putative tryptic phosphopeptide containing Thr9 would be very large and hydrophobic (and therefore probably insoluble) in 0.1% (v/v) trifluoracetic acid, the solvent used routinely to separate peptides by HPLC. In an initial experiment using unlabelled ATP, TIFA phosphorylated by ALPK1 was digested with chymotrypsin and subjected to mass spectrometry. A phosphopeptide was detected comprising amino acid residues 5-21 of TIFA phosphorylated at Thr9 (EDADT*EETVTCLQMTVY, where T* is Thr9). A further peptide was detected comprising residues 2-16 of TIFA phosphorylated at Thr9, which was preceded by the sequence Gly-Ser (GSTSFEDADT*EETVTCL, where T* is Thr9). The Gly-Ser sequence is derived from the linker region (GPLGS) between GST and TIFA that remains after cleavage of GST-TIFA with PreScission Protease. A third phosphopeptide was identified comprising the C-terminal peptide of TIFA, residues 167-184 (SLCSSQSSSPTEMDENES), but the site of phosphorylation could not be identified in this experiment (Fig 6F).

We next phosphorylated TIFA using Mg-γ^32^P-labelled ATP and subjected the chymotryptic digest to chromatography on a C18 column, which resolved five major ^32^P peptides C1-C5 (Fig 7A). Mass spectrometry indicated that peptides C1, C2 and C4 were all phosphorylated at Thr9 (Fig 7B). Solid phase sequencing of peptides C2 and C4 released ^32^P-radioactivity after the 7^th^ and 13^th^ cycles of Edman degradation, respectively, confirming that Thr9 was the site of phosphorylation (Fig 7C). The minor ^32^P-peptides C3 and C5 could not be identified by mass spectrometry.

**Figure 7.**
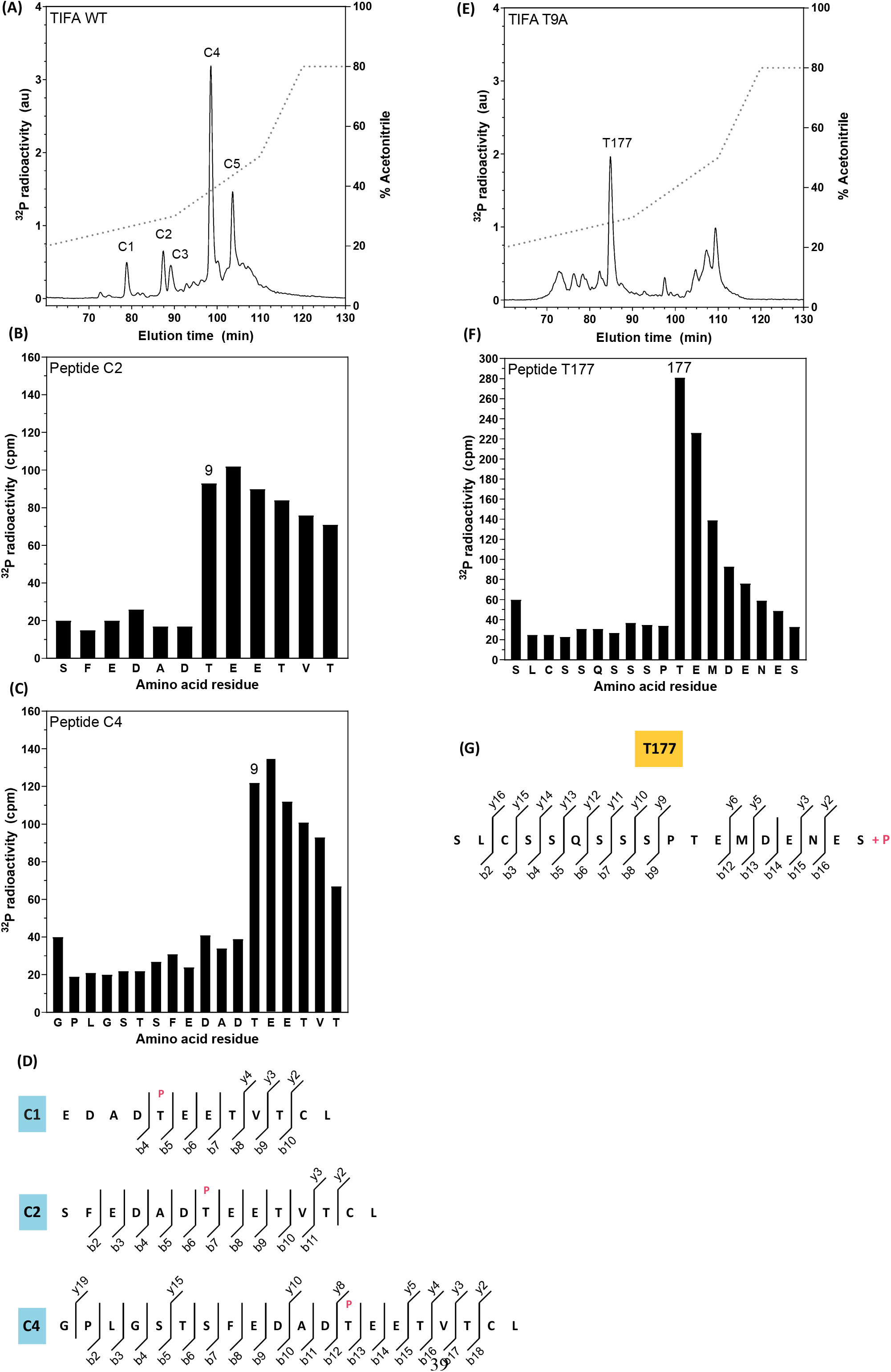
ALPK1 phosphorylates TIFA directly at Thr177 and Thr9. **(A)** ALPK1 KO HEK293 cells were re-transfected with plasmid DNA encoding FLAG-ALPK1 and FLAG-ALPK1 immunoprecipitated and used to phosphorylate TIFA as in Fig 6, except that the GST tag was first removed from GST-TIFA by cleavage with PreScission protease. The ^32^P-labelled band corresponding to TIFA was excised, digested for 24 h with chymotrypsin and the resulting peptides separated by HPLC with on-line radioactivity detection (see Methods). The solid line indicates the ^32^P-radioactivity and the broken line the acetonitrile gradient. **(B, C)** The peptides corresponding to C2 (**B**) and C4 (**C**) in **(A)** were subjected to solid phase sequencing and ^32^P-radioactivity released at each cycle of Edman degradation was quantitated by Cerenkov counting. **(D)** Peptides C1, C2 and C4 were subjected to mass spectrometry and the phospho-peptides detected are indicated along with their respective b- and y-ions, with the site of phosphorylation shown where localisation confidence was >90%. (**E**) As in (A) except that TIFA[T9A] was used at the substrate. **(F)** Peptide T177 from E was subjected to solid phase sequencing as in (B, C). (**G**) Peptide T177 was subjected to mass spectrometry and the fragmentation pattern observed in the mass spectrum is shown.

To identify the phosphorylated amino acid residue in the C-terminal peptide of TIFA, we phosphorylated TIFA[T9A] with ALPK1 and subjected it to more prolonged digestion with a 2-fold higher concentration of chymotrypsin. HPLC analysis of the digest revealed a major peptide, T177 (Fig 7D), with a molecular mass corresponding to amino acid residues 167-184 of TIFA (SLCSSQSSSPTEMDENES) plus one phosphate group (Fig 7E). Solid phase sequence analysis revealed that ^32^P-radioactivity was released from the peptide after the 11^th^ cycle of Edman degradation (Fig 7F), identifying Thr177 as the site of phosphorylation.

### Evidence that phosphorylation of Thr177 in the TRAF6-binding motif suppresses binding to TRAF6 but not TRAF2

Interestingly, Thr177 lies within a TRAF6-binding motif Pro-Xaa-Glu (Ye *et al*., 2002), where Xaa can be any amino acid residue. It has been reported that the TIFA[E178A] mutant, which disrupts the TRAF6 binding motif, fails to induce the oligomerisation and activation of TRAF6 in a cell-free system (Ea *et al*., 2004), and fails to stimulate NF-κB-dependent luciferase reporter gene expression when overexpressed in TIFA KO HEK293 cells (Zhou *et al*., 2018). Consistent with these findings, we found that ADP-heptose also failed to induce MAPK and IKK phosphorylation (Fig 8A) or interaction with TRAF6 (Fig 8B) when the TIFA[E178A] mutant was re-expressed in TIFA KO cells, in contrast to WT TIFA. However, the ADP-heptose-dependent interaction of TIFA[E178A] with TRAF2 and c-IAP1 was unaffected (Fig 8B). These findings imply that TRAF2 and TRAF6 bind to distinct sites on TIFA, and that TRAF2 is not captured by TIFA because it heterodimerises with TRAF6 (Middleton *et al*., 2017).

**Figure 8.**
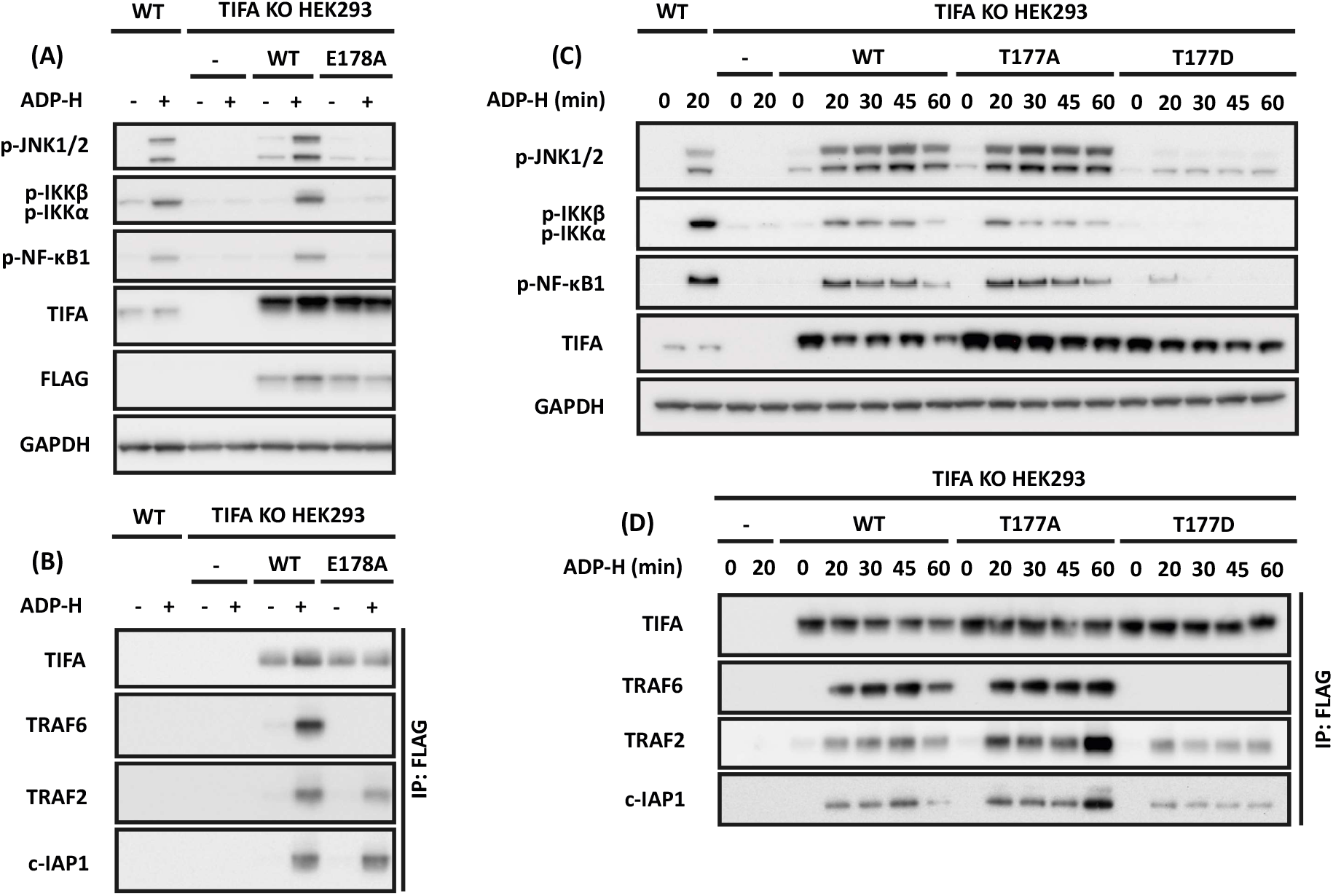
Mutations that disrupt the TRAF6-binding site of TIFA do not affect the interaction of TIFA with TRAF2. **(A)** TIFA KO HEK293 cells were re-transfected with 500 ng of plasmid DNA encoding empty vector (-), FLAG-TIFA or FLAG-TIFA[E178A]. 24 h later the cells were stimulated for 20 min with ADP-H and the extracts subjected to SDS-PAGE and immunoblotting with the antibodies indicated. **(B)** Cell extracts from (A) were immunoprecipitated with anti-FLAG and the immunoprecipitates analysed as in A. **(C)** As in A, except that plasmid DNA encoding empty vector, FLAG-TIFA, FLAG-TIFA[T177A] or FLAG-TIFA[T177D] was transfected **(D)** As in B but using extracts from C.

The TIFA[T177A] mutant did not impair ADP-heptose stimulated activation of MAPKs and IKKα/β or the interaction of TIFA with TRAF6, TRAF2 and c-IAP1 (Fig 8C and 8D). In contrast, and similar to the TIFA[E178A] mutant, the phospho-mimetic TIFA[T177D] mutant greatly reduced ADP heptose-stimulated MAPK and IKK phosphorylation (Fig 8C) and interaction with TRAF6 (Fig 8D), but the interaction with TRAF2 and c-IAP1 was unaffected (Fig 8D).

## Discussion

In this paper, we have made several findings that have advanced understanding of the ADP-heptose signalling network. First, we have established that the expression and activity of TAK1 is required for the ADP-heptose-dependent activation of the p38 MAP kinases, JNK1/2 and the canonical IKK complex. Second, that the K63-linked ubiquitin chains required to activate TAK1 are formed by both the TRAF6 and c-IAP1 E3 ligases, explaining why the E3 ligase activity of TRAF6 is not essential for the operation of the pathway. Third, we have found that ADP-heptose triggers the formation of M1-linked ubiquitin chains, which are known to facilitate activation of the canonical IKK complex.

Why the expression of TRAF6 is essential for the activation of MAP kinases and the canonical IKK complex, even though its E3 ligase activity is not, is unclear, but we speculate that the interaction of TRAF6 with TAK1, as well as the binding of K63-Ub chains to the TAB2/3 subunits of the TAK1 complex (Wang *et al*., 2001), may be needed for the activation of TAK1.

Although TRAF6 ubiquitylation was originally believed to be a key event in MyD88 signalling it was later reported that a TRAF6 mutant in which every Lys was mutated to Arg restored IL-1 signalling to TRAF6 KO fibroblasts (Walsh *et al*., 2008). Curiously, we now find that endogenous TRAF6 does not undergo ubiquitylation after stimulation with IL-1β in HEK293 cells or by R848 in BMDM. However, it does become ubiquitylated in response to stimulation with ADP-heptose. It will now be interesting to investigate whether TRAF6 ubiquitylation has an important role in ADP-heptose signalling.

The formation of M1-linked ubiquitin chains is catalysed by the HOIP component of LUBAC, which was recruited to the TIFA-TRAF6-TRAF2/c-IAP1 signalling complex together with its binding partner WRNIP1 in response to ADP-heptose (Fig 4A). An important role for M1-Ub chains in ADP-heptose signalling was indicated by the reduced ADP-heptose-dependent NFκB1 phosphorylation in MEFs expressing an E3 ligase-inactive mutant of HOIP (Fig S3D) or in MEFs expressing the NEMO[D311N] mutant that is unable to bind to M1-Ub linkages (Fig S3E).

A fourth finding that we made is that ADP-heptose stimulates the phosphorylation of the IKK-related kinase TBK1 at Ser172, the site required for TBK1 activation (Kishore *et al*., 2002). The ADP-heptose-stimulated phosphorylation of TBK1 was reduced in TRAF6 KO cells (Fig 3A) but did not require the expression or activity of TAK1 (Fig 2A) and was abolished in TRAF2/6 double KO cells (Fig S6). TRAF2 therefore has a role in the ADP-heptose-dependent activation of TBK1 that can be mediated independently of TRAF6 and does not simply function as a modulator of TRAF6-dependent signalling. An updated schematic of the ADP-heptose signalling pathway that incorporates our findings is shown in Fig 9.

**Figure 9:**
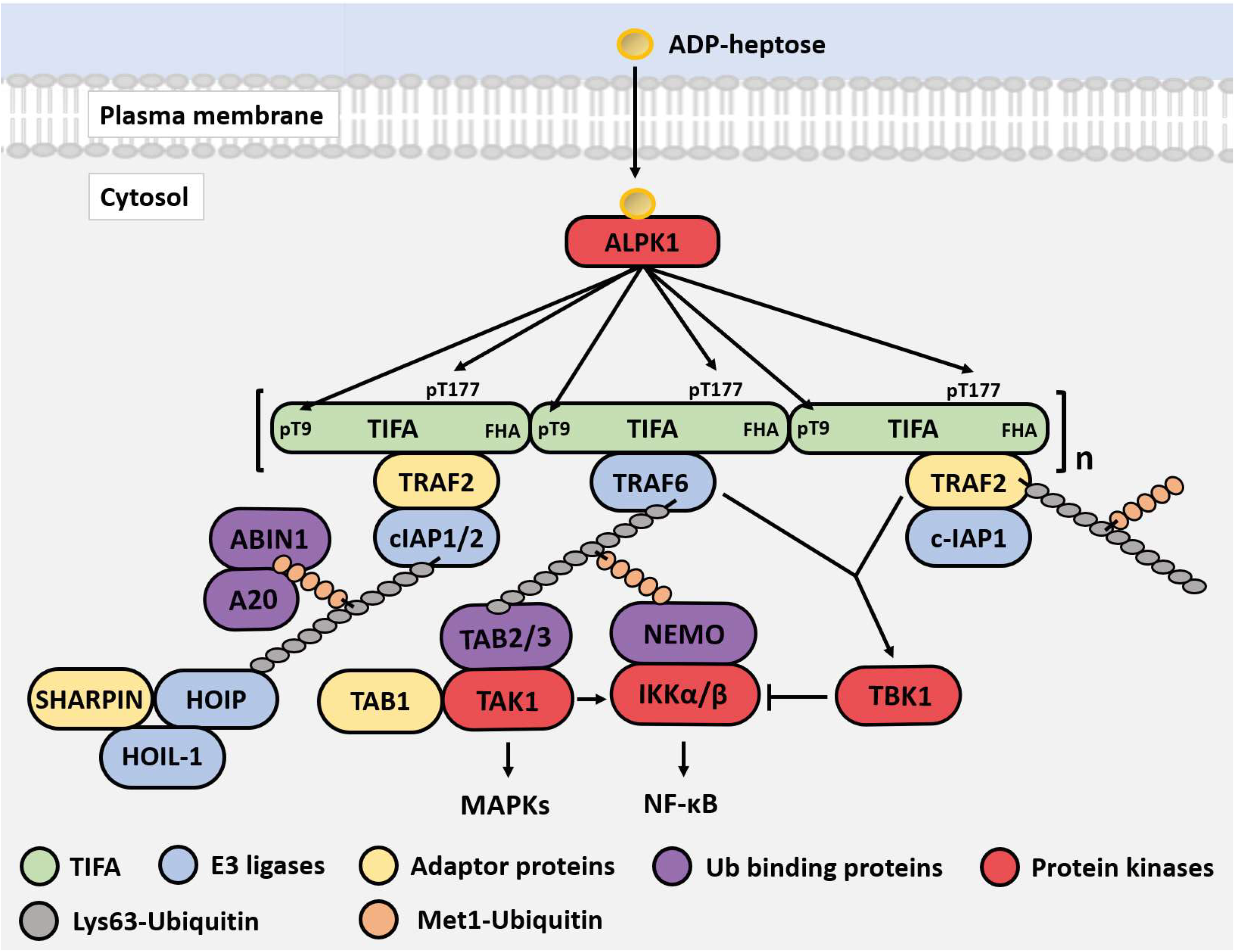
Schematic of the ADP-heptose signalling pathway in HEK293 cells. ADP-heptose enters the cytosol through an unidentified transporter where it binds to an allosteric site of ALPK1 inducing it’s activation and enabling it to phosphorylate TIFA at Thr9 and Thr177. The phosphorylated Thr9 interacts with the FHA domain of another TIFA molecule, leading to TIFA polymerisation in a head-to-tail fashion. This permits TRAF6 to interact with the TRAF6-binding motif of TIFA leading to the oligomerisation of TRAF6 and activation of its E3 ligase activity. The polymerised TIFA also recruits TRAF2 to an unknown site. The E3 ligase c-IAP1, which forms a complex with TRAF2, combines with TRAF6 to generate Lys63-linked ubiquitin oligomers that become attached covalently to TRAF6, TRAF2 and c-IAP1 and interact with the TAB2 or TAB3 components of TAK1 complexes, inducing the auto-activation of TAK1. ADP-heptose also stimulates the formation of Met1-linked ubiquitin chains that interact with NEMO to facilitate activation of the canonical IKK complex by TAK1. ADP-heptose additionally stimulates the activation of TBK1, which can be mediated by either TRAF6 or TRAF2 via an unknown pathway that does not require TAK1. TBK1 is likely to function as a negative feedback regulator of the pathway to prevent the overproduction of inflammatory mediators. The phosphorylation of TIFA at Thr177 also appears to be a feedback control device to suppress the TRAF6-dependent arm of the pathway. TBK1 may function at least in part, to restrict the activation of the canonical IKK complex (Clark *et al*., 2011). The phosphorylation of TIFA at Thr177 also appears to be a feedback control device to suppress the TRAF6-dependent arm of the pathway. By analogy with IL-1 and TLR signalling (Emmerich *et al*., 2013), the K63-Ub and M1-Ub chains are depicted as being linked to one another, although this has not yet been established.

One of the physiological roles of TBK1 is to restrict the activation of the canonical IKK complex (Clark *et al*., 2011) by phosphorylating inhibitory sites on IKKα/β and on their regulatory subunit NEMO (Delhase *et al*., 1999; Palkowitsch *et al*., 2008). Such mechanisms are needed to prevent the hyperactivation of innate immune signalling pathways, which would otherwise lead to the overproduction of inflammatory mediators that cause inflammatory and autoimmune diseases. It is therefore noteworthy that two other negative regulators of innate immune signalling pathways, the ubiquitin-binding proteins A20 and ABIN1 (A20-binding inhibitor of NF-κB1) (reviewed (Cohen and Strickson, 2017)), were also recruited to the ADP-heptose-stimulated signalling complex (Fig 4A).

We have also established for the first time that ALPK1 phosphorylates TIFA directly at Thr9 and identified Thr177 as a second site phosphorylated by ALPK1. Interestingly, Thr177 lies in the TRAF6-binding motif of TRAF6 (Pro-Thr-Glu) and its mutation to Asp to try and mimic the effect of phosphorylation by introducing a negative charge at this position, suppressed the binding of TRAF6 to TIFA and consequently abolished the ADP-heptose-stimulated phosphorylation of MAP kinases and the canonical IKK complex (Fig 8). The ALPK1-catalysed phosphorylation of TIFA at Thr177 may represent yet a further mechanism for restricting the strength of activation of the ADP-heptose signalling pathway. The TIFA[T177D] mutation did not affect interaction with TRAF2, indicating that the binding of TRAF2 to TIFA occurs independently of the binding of TRAF6. The TIFA[T9A] and other TIFA mutants prevent the binding of both TRAF6 and TRAF2 because they prevent the formation of TIFA oligomers, which is essential to recruit the proteins needed to propagate the ADP-heptose signal. TIFA does not possess either of the known TRAF2-binding motifs Pro/Ser/Ala/Thr-Xaa-Gln/Glu-Glu and Pro-Xaa-Gln-Xaa-Xaa-Asp (Ye *et al*., 1999), and it will be important to define the TRAF2-binding site in TIFA in future studies.

## Materials and Methods

### Antibodies

The following antibodies were obtained from Cell Signalling Technology (CST): phospho-specific antibodies recognising IKKα phosphorylated at Ser176 and Ser180 and IKKβ phosphorylated at Ser177 and Ser181 (#2697), p38α MAPK and p38γ MAPK phosphorylated at Thr180 and Tyr182 within their Thr-Gly-Tyr motifs (#9211), JNK1 and JNK2 phosphorylated at Thr183 and Thr185 within their Thr-Pro-Tyr motifs (#9251), TAK1 phosphorylated at Thr187 (#4536), TBK1 phosphorylated at Ser172 (#5483) and IKKε phosphorylated at Ser172 (#8766) (these are the phosphorylation sites within the activation loops that trigger the activation of these protein kinases); the IKKβ substrate NF-κB1 (also called p105) phosphorylated at Ser133 (#4806). Also from CST were antibodies recognising anti-rabbit IgG (#7074) and anti-mouse IgG (#7076), FLAG (#2368), GAPDH (#2118), human c-IAP1 (#7065), human c-IAP2 (#3130), IκBα (#4812), IKKβ (#8943), IRAK1(#4504), NIK (#4994), Pellino 1 (#31474), human TIFA (#61358), TRAF2 (#4712) and human TRAF6 (#8028). An Antibody recognising anti-human IgG (#2040-05) was from Southern Biotech and anti-rat IgG (#405405) from Biolegend. Anti-sheep IgG (#ab97130) and murine TRAF6 (#ab40675) were from Abcam and murine c-IAP1 (#803-335-C100) was from Enzo Life Sciences. Antibodies recognising Met1-linked ubiquitin and Lys63-linked ubiquitin were generous gifts from Vishva Dixit (Genentech). Antibodies against ALPK1[1-358] (S593D, bleed 5) and ALPK1[395-823] (S594D, bleed 3) were raised in sheep by the MRC Reagents and Services section of the MRC Protein Phosphorylation and Ubiquitylation Unit, University of Dundee and are available on request (mrcppureagents.dundee.ac.uk).

### DNA constructs

The following DNA vectors for expression in mammalian cells were made by the DNA cloning team of MRC Reagents and Services, MRC Protein Phosphorylation and Ubiquitylation Unit, University of Dundee and are available on request (mrcppureagents.dundee.ac.uk). For the generation of stable cell lines using pBabe and pRetroXTight vectors: TRAF6 (DU47223), FLAG-TRAF6 (DU32495), TRAF6[L74H] (DU47224), TRAF6[120-522] (DU51445), TAK1 (DU51270), and TAK1 [D175A] (DU51293). For transient transfection: FLAG-ALPK1 (DU65668, CMV promoter), FLAG-ALPK1 (DU65976, UbC promoter), FLAG-ALPK1[K1067M] (DU65680, CMV promoter), FLAG-ALPK1[K1067M] (DU71020, UbC promoter), FLAG-TIFA (DU71017, UbC promoter), FLAG-TIFA[T9A] (DU71055, UbC promoter), FLAG-TIFA[T9D] (DU71053), FLAG-TIFA[T9E] (DU71054, UbC promoter), FLAG-TIFA[T9S] (DU71067, UbC promoter), FLAG-TIFA[T177A] (DU71049, UbC promoter), FLAG-TIFA[T177D] (DU71051, UbC promoter), and FLAG-TIFA[E178A] (DU71057, UbC promoter). The plasmids with a CMV promoter were used for purification of proteins, whereas those with the weaker UbC promoter (Qin *et al*., 2010) were used for the study of intracellular signalling.

The procedure and guide sequences for the removal of TAK1 or TRAF6 are described elsewhere (Strickson *et al*., 2017; Zhang *et al*., 2017). The guides for removal of TRAF2 were GACCCTCCTGGGGACCAAGC (Sense, DU52594) and GCCGGGCTGTAGCAACTCCA (Antisense, DU52604).

### Proteins

Glutathione-S-transferase (GST)-tagged proteins contained a PreScission protease cleavage site between the GST and the protein of interest: GST-TIFA (DU2421), GST-TIFA[T9A] (DU65850), and GST-TIFA[T9S] (DU65858). These proteins were produced by the MRC Protein Phosphorylation and Ubiquitylation Unit’s reagents section and are available on request (mrcppureagents.dundee.ac.uk). Halo-NEMO (DU35939) was expressed and coupled to Halo-link resin as described (Emmerich *et al*., 2013). The protein phosphatase from bacteriophage λgt10 (Phage λ phosphatase) was purchased from New England Biolabs.

### Cell culture, cell maintenance and cell lysis

HEK293 cells stably expressing relatively low levels of the IL-1 receptor, termed IL-1R* HEK293 cells, has been described (Strickson *et al*., 2017). ALPK1 knockout and TIFA knockout HEK293 cells and their parental cell line were purchased from Invivogen and termed HEK293 cells. HEK293 cells were cultured at 37°C in a humidified atmosphere containing 5% CO2. The growth media contained Dulbecco’s modified Eagle’s medium (DMEM), containing 25 mM HEPES pH 7.3, 10% (v/v) foetal bovine serum (FBS), 2 mM L-glutamine, 100 U/ml penicillin and 0.1 mg/ml streptomycin. BMDM were obtained by differentiating bone marrow extracted from the femur and tibia of mice and cultured as described (Pauls *et al*., 2013). Since few TRAF6 KO mice are born alive and those that do die within days, macrophages from TRAF6 KO mice were produced from foetal liver cells and cultured as described (Strickson *et al*., 2017). The preparation of MEFs from HOIP[C879S] and NEMO[D311N] mice has been described previously and were cultured as described (Zhang *et al*., 2014). Cells were washed twice with ice-cold PBS, scraped from the plate in lysis buffer (50 mM Tris/HCl pH 7.5, 1 mM EDTA, 1 mM EGTA, 1% (v/v) Triton X-100, 1 mM sodium orthovanadate, 50 mM sodium fluoride, 5 mM sodium pyrophosphate, 270 mM sucrose, 10 mM sodium 2-glycerophosphate, 0.2 mM phenylmethylsulfonyl fluoride and 1 mM benzamidine, supplemented with complete protease inhibitor cocktail). When studying ubiquitin chain formation, 100 mM iodoacetamide was included to inhibit deubiquitylases (DUBs) (Emmerich and Cohen, 2015). Cell lysates were clarified by centrifugation for 20 min at 20,000 x *g* at 4 °C and the supernatants (cell extracts) transferred to 1.5 ml microcentrifuge tubes, snap frozen and stored at -80 °C. The protein concentrations of cell extracts were determined by the Bradford procedure.

### Transient transfection of HEK293 cells

HEK293 cells were transfected in 10 cm dishes at 70% confluency using lipofectamine 2000 according to the manufacturer’s instructions.

### Generation of retroviruses using HEK293FT cells and infection of target cells

HEK293FT cells were plated on 10 cm dishes pre-coated with poly-L-lysine. Transfection was performed at 90% confluency using Lipofectamine LTX according to the manufacturer’s instructions. Retroviral particles were prepared using pBabe and pRetroXTight vectors as described elsewhere (Strickson *et al*., 2017; Zhang *et al*., 2017). HEK293 cells in 6-well plates were infected at 50% confluency using 500 µl of viral suspension and 8 µg/ml polybrene. The media was replaced 24 h later and after a further 24 h, the retrovirally transduced cells were selected for stable integration of the protein of interest using media containing 2 µg/ml puromycin and/or 1 mg/ml G418.

### Mice

TRAF6[L74H] knock-in mice were produced as described (Strickson *et al*., 2017) and heterozygous TRAF6 KO mice were provided by Tak Mak. Mice were maintained on a C57BL/6 background and provided with free access to food and water. Animals were kept in ventilated cages under specific pathogen-free conditions in accordance with UK and European Union regulations. Experiments were carried out subject to approval by the University of Dundee ethical review board under a UK Home Office project licence.

### Agonists and inhibitors

ADP-D,D-heptose and ADP-L,D-heptose was synthesised as described (Zamyatina *et al*., 2003) and were free of endotoxin contamination as judged by the QCLTM Kinetic Chromogenic LAL Assay (Lonza). The ADP-heptoses, Resiquimod (R848) (Invivogen) and human IL-1β (Invivogen), were dissolved in PBS and added to the cell culture medium to achieve final concentrations of 10 μM (ADP-heptose), 250 ng/ml (R848), and 5 ng/ml (IL-1β). NG-25, BV-6 and Birinapant (from SelleckChem) and (5Z)-7-Oxozeaenol (from Tocris) were dissolved in DMSO at 10 mM and added to achieve the final concentrations indicated in figure legends. An equivalent volume of DMSO was added to the culture medium in control incubations.

### SDS-PAGE, transfer to PVDF membranes and immunoblotting

Reduced cell extracts (20 µg of protein) or immunoprecipitates were resolved by SDS-PAGE and transferred to PVDF membranes (Merck-Millipore, #10344661) using a Trans-Blot transfer cell (BioRad). The membranes were blocked by incubation for 1 h in 50 mM Tris/HCl pH 7.5, 150 mM NaCl, and 0.1% (v/v) Tween-20 (TBS-T) containing 5% (w/v) non-fat milk powder (Marvel). Primary antibody in TBS-T containing 5% (w/v) bovine serum albumin was then incubated with the PVDF membrane on a shaker for 16 h at 4 °C. The immunoblots were washed five times (10 min per wash) with TBS-T followed by incubation for 1 h with the appropriate horseradish peroxidase-conjugated secondary antibody in TBS-T containing 5% (w/v) non-fat milk powder and washed a further five times with TBS-T. Most blots were developed by incubation at ambient temperature with Amersham ECL Select Western Blotting Detection Reagent (GE Healthcare) or Super-Signal West Pico Chemiluminescent Substrate (Merck-Millipore). The most sensitive reagent, Clarity Max Western ECL Substrate (Bio-Rad), was used to blot for ubiquitylated proteins. Most immunoblots were imaged using a ChemiDoc MP scanner (#17001402, Bio-Rad). Ubiquitin blots were exposed to Amersham Hyperfilm X-ray film in an X-ray cassette and developed using a Konica automatic developer. Antibodies were stripped from membranes by incubation for 15 min at ambient temperature with Restore Western Blot Stripping Buffer (Thermo Fisher). The stripped membranes were washed twice with TBS-T and immunoblotting repeated by incubation with a different primary antibody.

### Immunoprecipitation of endogenous ALPK1

Cell extracts (1 mg protein) were treated for 15 min at ambient temperature with benzonase (Merck-Millipore) and pre-cleared by incubation for 30 min at 4°C with 20 µl of protein A/G-Sepharose beads. ALPK[1-358] antibody (10 µg) was added to the supernatant and incubated for 1 h at 4°C on a rotating wheel followed by addition of 20 µl protein A/G-Sepharose for 30 min. After brief centrifugation, the supernatant was discarded and the beads washed twice with 50 mM Tris-HCl pH 7.5, 1% (v/v) Triton X-100 and 500 mM NaCl and then twice with 50 mM Tris-HCl pH 7.5, 1% (v/v) Triton X-100. The immunoprecipitated ALPK1 was released by incubation for 5 min at 95 °C with LDS sample buffer (Thermo Fisher, #NP0007) containing 2.5% (v/v) β-mercaptoethanol. The suspension was transferred to Spin-X columns (Corning, #8162), centrifuged for 1 min at 20,000 x *g* and the flowthrough was analysed by SDS-PAGE and immunoblotted with an antibody recognising ALPK1[395-823].

### Halo-NEMO pulldowns

Packed Halo-NEMO resin (20 µl) was added to cell extracts (2 mg protein) and incubated for 4 h at 4 °C on a rotating wheel to capture Met1/Lys63-linked ubiquitin chains and the proteins to which they were attached covalently and non-covalently. The resin was washed twice with 50 mM Tris-HCl pH 7.5, 1% (v/v) Triton X-100 and 500 mM NaCl and twice with 50 mM Tris-HCl pH 7.5, 1% (v/v) Triton X-100 to remove proteins attached non-specifically to the resin. Proteins captured on the resin were released by incubation for 10 min at 37 °C in LDS sample buffer containing 2.5% (v/v) β-mercaptoethanol. The suspension was transferred to Spin-X columns (Corning, #8162), centrifuged for 1 min at 20,000 x *g* and the flowthrough was analysed by SDS-PAGE.

### Immunoprecipitation of FLAG-tagged proteins

To identify proteins interacting with TRAF6 by MS, HEK293 cells stably expressing FLAG-TRAF6 were either left untreated, stimulated for 15 min with IL-1β or ADP-heptose. Cells were lysed in ice-cold lysis buffer containing 100 mM iodoacetamide, centrifuged (30 min at 20,000 x *g* at 4 °C) and the cell extracts passed through 0.45 µm filters. The supernatants from unstimulated, IL-1β- and ADP-heptose-stimulated cells (30 mg protein per condition) were diluted to 2 mg/ml protein with lysis buffer and pre-cleared by incubation for 12 h at 4°C on a rotating wheel with mouse IgG-agarose. The supernatants were collected and incubated for 4 h at 4 °C on a rotating wheel with 40 μl of packed anti-FLAG M2 affinity gel (Sigma, #A2220), which had been washed three times with lysis buffer prior to use. After centrifugation, the supernatant was discarded and the pelleted gel washed three times with 50 mM Tris-HCl pH 7.5, 1% (v/v) Triton X-100 containing 300 mM NaCl and twice with 50 mM Tris-HCl (pH 7.5), 1% (v/v) Triton X-100. The proteins captured on the anti-FLAG resin were eluted by incubation with 46 μl of TBS containing 150 ng/µL of 3X FLAG peptide (Sigma, #F4799). For mass spectrometry (MS) analysis the above procedure was repeated 5 times for each of the three conditions, giving a total of 15 samples.

FLAG-ALPK1 or FLAG-TIFA were transiently re-expressed in ALPK1 KO or TIFA KO HEK293 cells, respectively, for 48 h unless stated otherwise and the proteins were immunoprecipitated with anti-FLAG and washed as described above, except that 0.2-1 mg of cell extract protein and 20 µl of resin and elution from the anti-FLAG gel was achieved using LDS sample buffer as described earlier. To identify sites on TIFA phosphorylated by FLAG-ALPK1, the FLAG-ALPK1 was immunoprecipitated as described above but washed with 500 mM instead of 300 mM NaCl.

### Phosphorylation of TIFA by ALPK1 in vitro

GST-TIFA was phosphorylated using FLAG-ALPK1 attached to the anti-FLAG resin and carried out for 20 min in 25 µl of 50 mM Tris/HCl pH 7.5, 2 mM DTT, 0.1 mM EGTA, 8 μM GST-TIFA, 10 nM ADP-heptose, 10 mM magnesium acetate and 0.1 mM [γ-^32^P]ATP (specific radioactivity 1000 cpm/pmol). Reactions were terminated using LDS sample buffer containing 2.5% (v/v) β-mercaptoethanol and heated for 5 min at 75 °C. The resin was pelleted by brief centrifugation at 13,000 x *g* and half of the supernatant subjected to SDS-PAGE. After staining for 1 h with Instant Blue and destaining in water for 48 h, incorporation of ^32^P-radioactivity into TIFA and ALPK1 was analysed by autoradiography and Cerenkov counting.

### Identification of phosphorylation sites in TIFA

To identify the sites on TIFA phosphorylated by FLAG-ALPK1, GST-TIFA and GST-TIFA[T9A] were cleaved with PreScission protease, and passed through glutathione-Sepharose to remove GST, prior to phosphorylation. The purified TIFA preparations began with linker sequence GLPGS followed by amino acid residues 2-184 of TIFA. Following phosphorylation for 20 min, the ^32^P-labelled band corresponding to phosphorylated TIFA was excised, cut into many sections and washed with 50 mM Tris-HCl pH 8.0 containing 50% (v/v) acetonitrile until the gel pieces were colourless as described previously (Shevchenko *et al*., 2007). The gel pieces were then incubated for 30 min at ambient temperature with 100 mM triethyl ammonium bicarbonate (TEAB) containing 10 mM dithiothreitol and alkylated in the dark for 30 min at ambient temperature in 50 mM iodoacetamide. The gel pieces were resuspended in 0.1 ml of 100 mM Tris/HCl pH 8.0 containing 10 mM calcium chloride and chymotrypsin (Promega, #V1061) for 16 h at 25 °C. The resulting peptides were collected and separated on a reverse-phase Vydac C18 column (Separations Group, CA, USA) with on-line radioactivity detection. The column was equilibrated in 0.1% (v/v) trifluoroacetic acid and developed with a linear acetonitrile gradient at a flow rate of 0.2 ml/min. Fractions (0.1 ml) were collected and analysed for ^32^P radioactivity by Cerenkov counting. The identified phosphopeptide fractions were analysed by liquid chromatography (LC)-MS/MS using a Thermo U3000 RSLC nano liquid chromatography (LC) system or an EvoSep LC coupled to an Exploris 240 mass spectrometer (Thermo Fisher) to determine the primary sequence of the phosphopeptides. Data files were searched using Mascot (www.matrixscience.com) with a 10 ppm mass accuracy for precursor ions, a 0.06 Da tolerance for fragment ions, and allowing for Phospho (ST), Phospho (Y), and oxidation of methionine (M) as variable modifications. Individual MS/MS spectra were inspected using Xcalibur 2.2 and Proteome Discoverer with phosphoRS 3.1 (Thermo Fisher) was used to assist the assignment of phosphorylation sites. The sites of phosphorylation of ^32^P-labeled peptides were also determined by solid-phase Edman degradation on a Shimadzu PPSQ33A Sequencer (Kyoto, Japan) with the peptides coupled to an 80 Sequelon-AA membrane (Applied Biosystems), as described previously (Campbell and Morrice, 2002).

### TMT labelling and mass spectrometry

The Flag-TRAF6 preparations eluted from FLAG resin with 3X FLAG peptide were digested with a combination of trypsin and Lys-C protease (Promega, #V5071) using S-trap mini columns (Protifi) according to the manufacturer’s instructions, except that 20 mM iodoacetamide was used for alkylation and washing steps with 100 mM TEAB in 90% (v/v) methanol repeated 10 times to remove traces of Tris buffer that would otherwise interfere with TMT labelling. The resultant peptides were resuspended for 15-plex TMT labelling according to the manufacturer’s protocol (Thermo Fisher) and the 15 samples (see the preceding section) were combined in equimolar amounts and dried. Peptides were resuspended in 100 μl of 5 mM ammonium acetate at pH 10 and injected on to a XBridge Peptide BEH C18 column (Waters #186003561). Peptides were eluted from the column using basic reverse phase fractionation (using 10 mM ammonium acetate in water as Buffer A and 10 mM ammonium acetate in 80:20 (v/v) acetonitrile:water as Buffer B) on a 55 min multistep gradient at 100 μl/min. Eluted peptides were collected into 96 fractions, pooled into 24 fractions using non-consecutive concatenation and dried. The peptides were redissolved in 5% (v/v) formic acid and injected on a UltiMate 3000 RSLCnano System coupled to a Orbitrap Fusion Lumos Tribrid Mass Spectrometer (Thermo Fisher). Peptides were loaded on to an Acclaim Pepmap trap column (Thermo Fisher #164564-CMD) prior to analysis on a PepMap RSLC C18 analytical column (Thermo Fisher #ES903) and eluted using a 120 min linear gradient from 3 to 35% Buffer B (Buffer A: 0.1% formic acid, Buffer B: 0.08% formic acid in 80:20 (v/v) acetonitrile:water. The eluted peptides were then analysed by the mass spectrometer operating in synchronous precursor selection mode (SPS) on a TOP 3s method.

### Analysis of TMT MS data

A peptide search was conducted by interrogating the Uniprot Swissprot Human database (released on 05/10/2021) using MaxQuant 1.6.17.0 in MS3 reporter ion mode. using default parameters with the addition of deamidation (NQ) and phospho (STY) as variable modifications. Protein groups present in the reversed database, or a database of proteins that are frequent contaminants, or proteins in which only one unique peptide or razor peptide were detected, or proteins present in less than 4 of the 5 replicates were excluded. Missing values were then imputed using a Gaussian distribution centred on the median with a downshift of 1.8 and width of 0.3 (based on the standard deviation) and protein intensities were median normalized. Protein regulation was assessed using a 2 sample Welch test and P-values were adjusted using Benjamini Hochberg (BH) multiple hypothesis correction. Proteins in TRAF6 immunoprecipitates were considered-to-be enriched significantly after cell stimulation with ADP-heptose if the BH corrected P-value was smaller than 0.05 and the fold-change was greater than 1.75. The mass spectrometry proteomics data have been deposited to the ProteomeXchange Consortium via the PRIDE (Perez-Riverol *et al*., 2022) partner repository with the dataset identifier PXD034964.

## Acknowledgements

We thank Nicola Wood and Thomas Macartney (MRC-PPU Reagents and Services) for making the DNA constructs and CRISPR reagents, respectively, and Clara Figueras Vadillo for genotyping and maintenance of mice and the differentiation of bone marrow and foetal liver into macrophages. T.S. acknowledges the award of a Ph.D. studentship 2087974 from the UK Medical Research Council (MRC). The study was also supported by Wellcome Trust Investigator Award 209380/Z/17/Z and by MRC Programme Grant MR/R021406/1 (to P.C.).

## Author contributions

The project was conceptualised by T.S and P.C, who also wrote the manuscript. N.S. synthesised ADP-heptoses, R.G. performed HPLC, Edman sequencing and mass spectrometry on samples prepared by T.S. relating to Fig 7 and F.L. performed mass spectrometry on samples prepared by T.S. and analysed data relating to Fig 4A. All other experiments were performed and analysed by T.S, who also prepared the figures.

## Supplementary Figures

**Figure S1.**
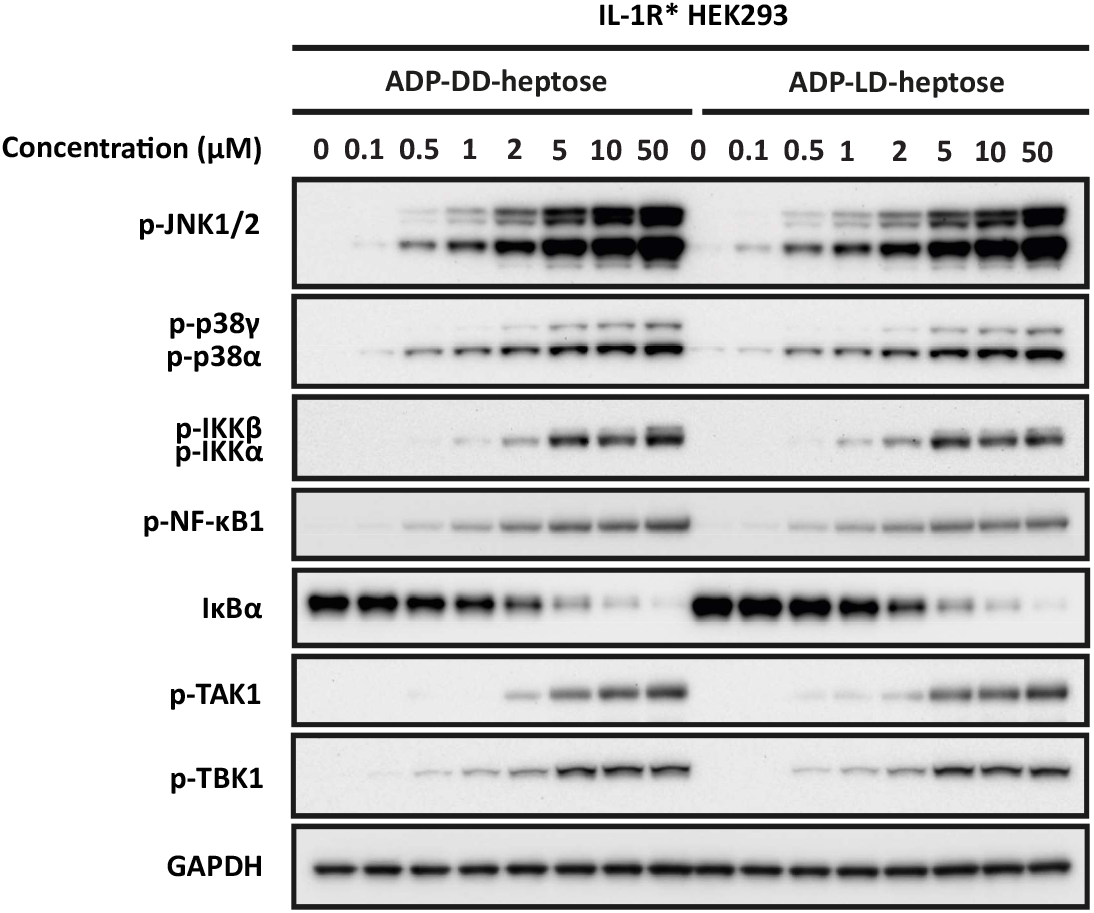
Concentration of ADP-DD-heptose and ADP-LD-heptose required for maximal activation of MAP kinases and the canonical IKK complex. IL-1R* HEK293 cells were stimulated for 30 min with the indicated concentrations of ADP-DD-heptose or ADP-LD-heptose. Cell extracts were analysed by SDS-PAGE and immunoblotted with antibodies recognising the proteins indicated.

**Figure S2.**
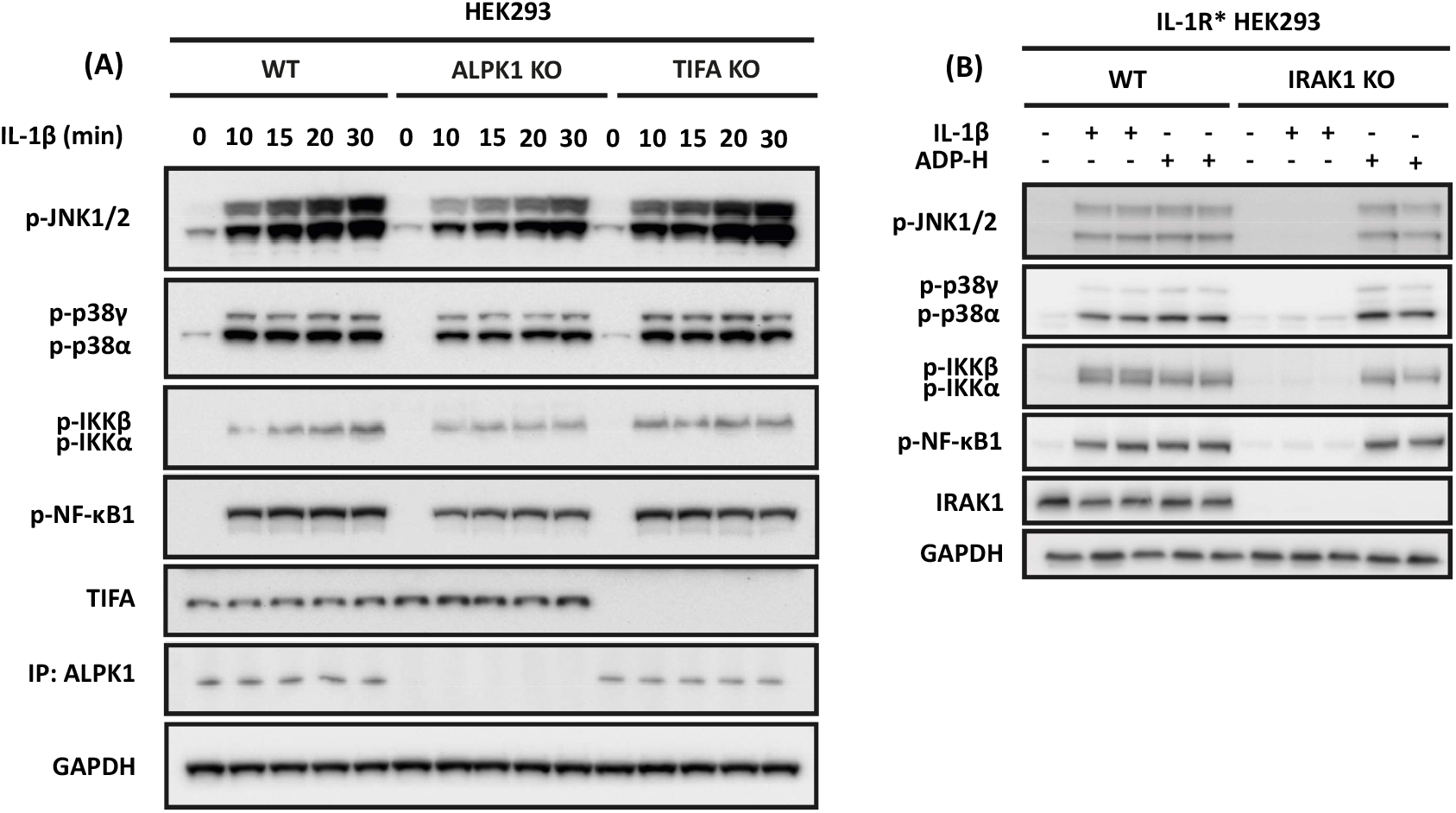
ALPK1 and TIFA are not required for IL-1β signalling and IRAK1 is not required for ADP-heptose signalling. **(A)** Parental, ALPK1 KO and TIFA KO HEK293 cells were stimulated with IL-1β the times indicated. Cell extracts were then analysed by SDS-PAGE and immunoblotting with the antibodies indicated. ALPK1 was immunoblotted after first immunoprecipitating it from the cell extracts. (B) As in A, except that parental and IRAK1 KO IL-1R* HEK293 cells were stimulated for 20 min with ADP-H or IL-1β.

**Figure S3.**
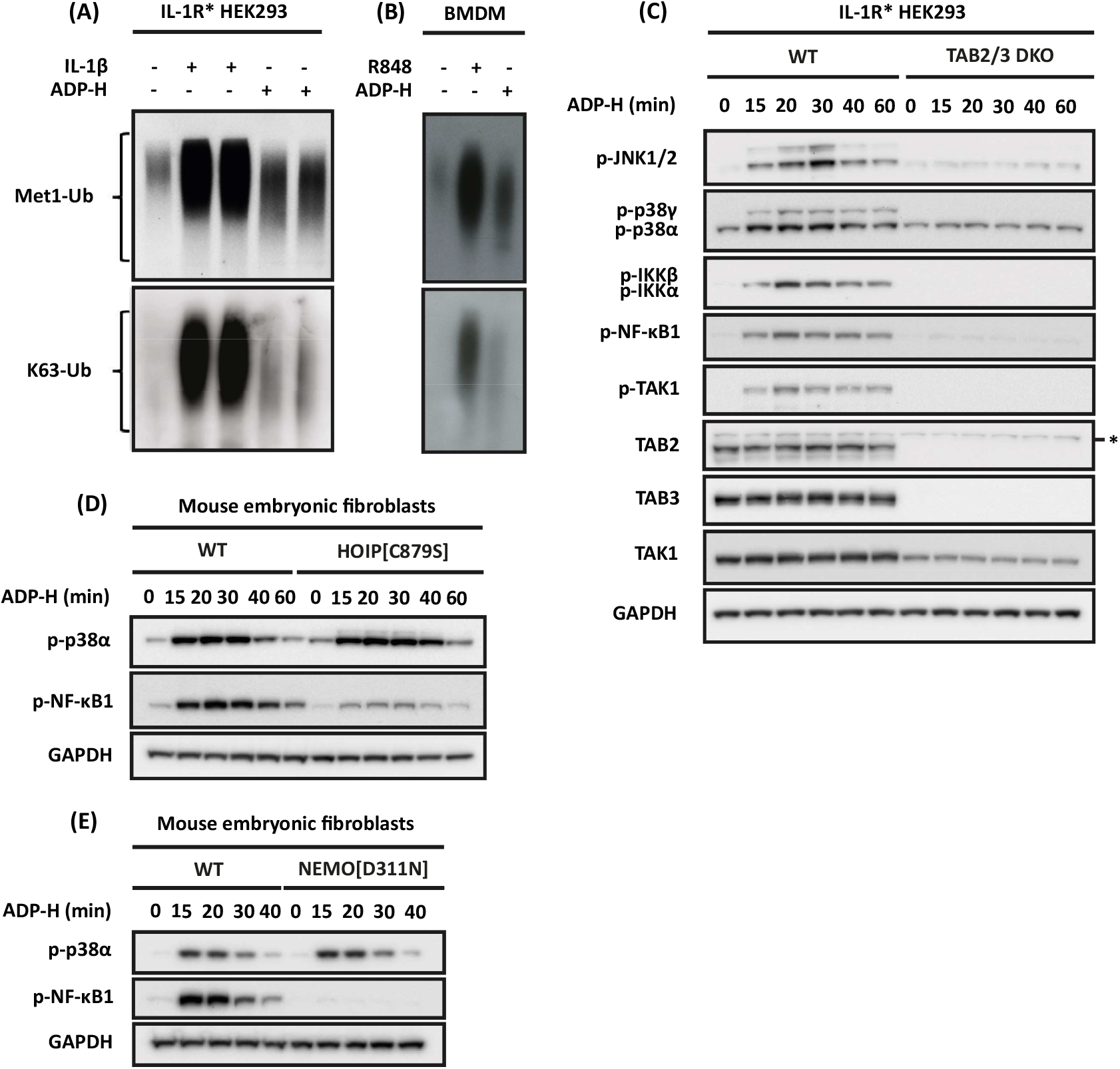
ADP-heptose signalling induces the formation of Lys63-linked and Met1-linked ubiquitin chains and requires the expression of TAB2 and TAB3, the E3 ligase activity of HOIP and the interaction of ubiquitin chains with NEMO. **(A)** IL-1R* HEK293 cells were stimulated for 15 min with ADP-heptose (ADP-H) or IL-1β. Ubiquitin chains were captured from the cell extracts on Halo-NEMO beads (see methods) and detected by SDS-PAGE and immunoblotting with antibodies recognising K63- or M1-linked ubiquitin oligomers. **(B)** As in A, but using primary BMDM from WT mice stimulated for 15 min with ADP-H or R848. (**C**) Parental and TAB2/3 double KO IL-1R* HEK293 cells were stimulated with ADP-H for the times indicated and immunoblotting of the cell extracts performed with the antibodies indicated. An asterisk indicates protein(s) recognised non-specifically by an antibody. **(D, E)** MEFs from WT and HOIP[C879S] (**D**) or WT and NEMO[D311N] **(E)** mice were stimulated for the times indicated with ADP-H and immunoblotting of the cell extracts performed with the antibodies indicated.

**Figure S4.**
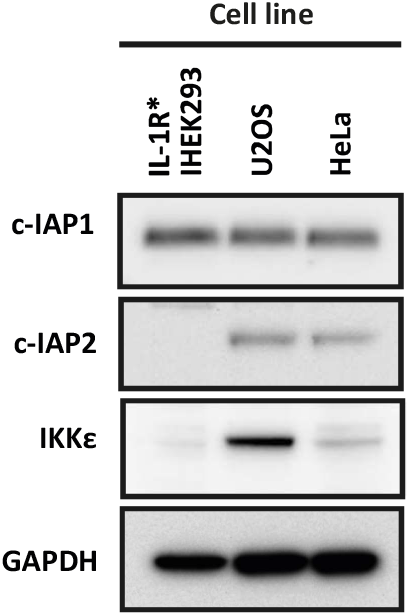
Relative levels of expression of c-IAP1, c-IAP2 and IKKε in three human cell lines. Cell extracts from three human cell lines were analysed by SDS-PAGE and immunoblotted with antibodies recognising the proteins indicated.

**Figure S5.**
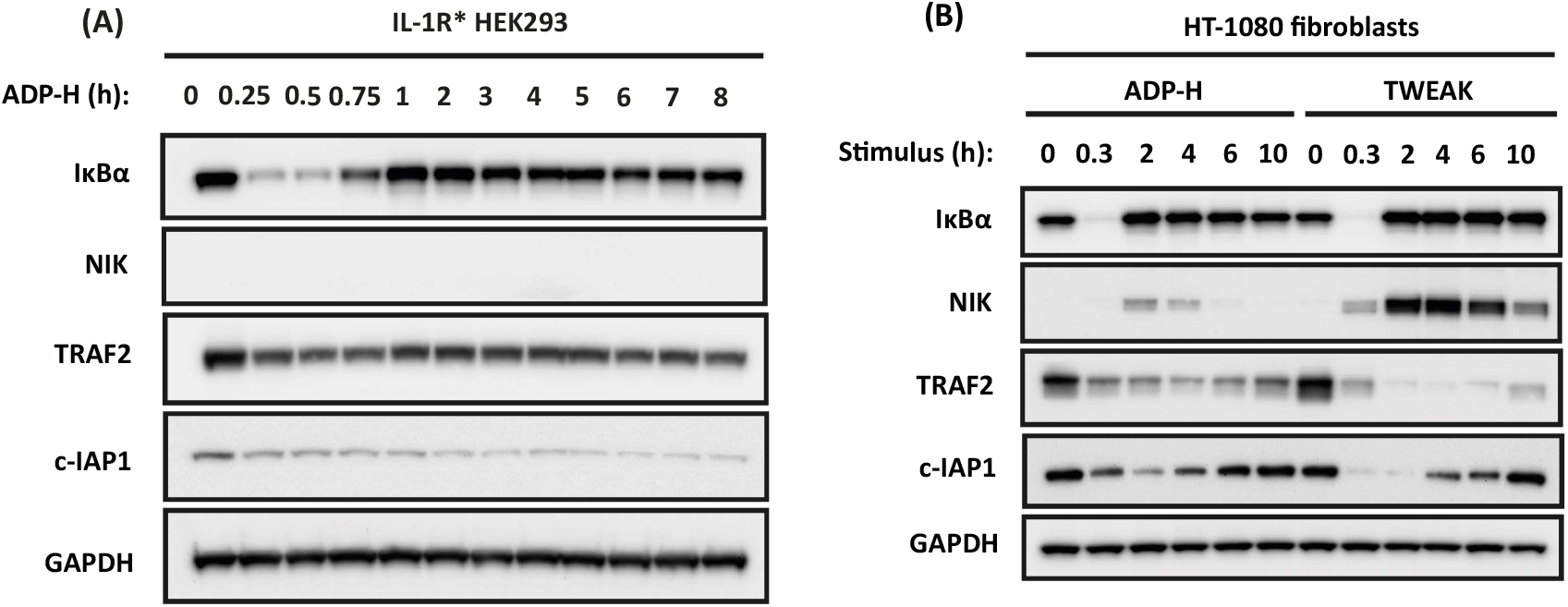
ADP-heptose does not activate the non-canonical NF-κB signalling pathway in HEK293 cells but activates it weakly in HT-1080 fibroblasts. **(A)** IL-1R* HEK293 cells were stimulated for the times indicated times with ADP-heptose (ADP-H) and the cell extracts analysed by SDS-PAGE and immunoblotting with antibodies recognising the proteins indicated **(B)** As in A except that HT-1080 cells were stimulated with ADP-H or TWEAK.

**Figure S6.**
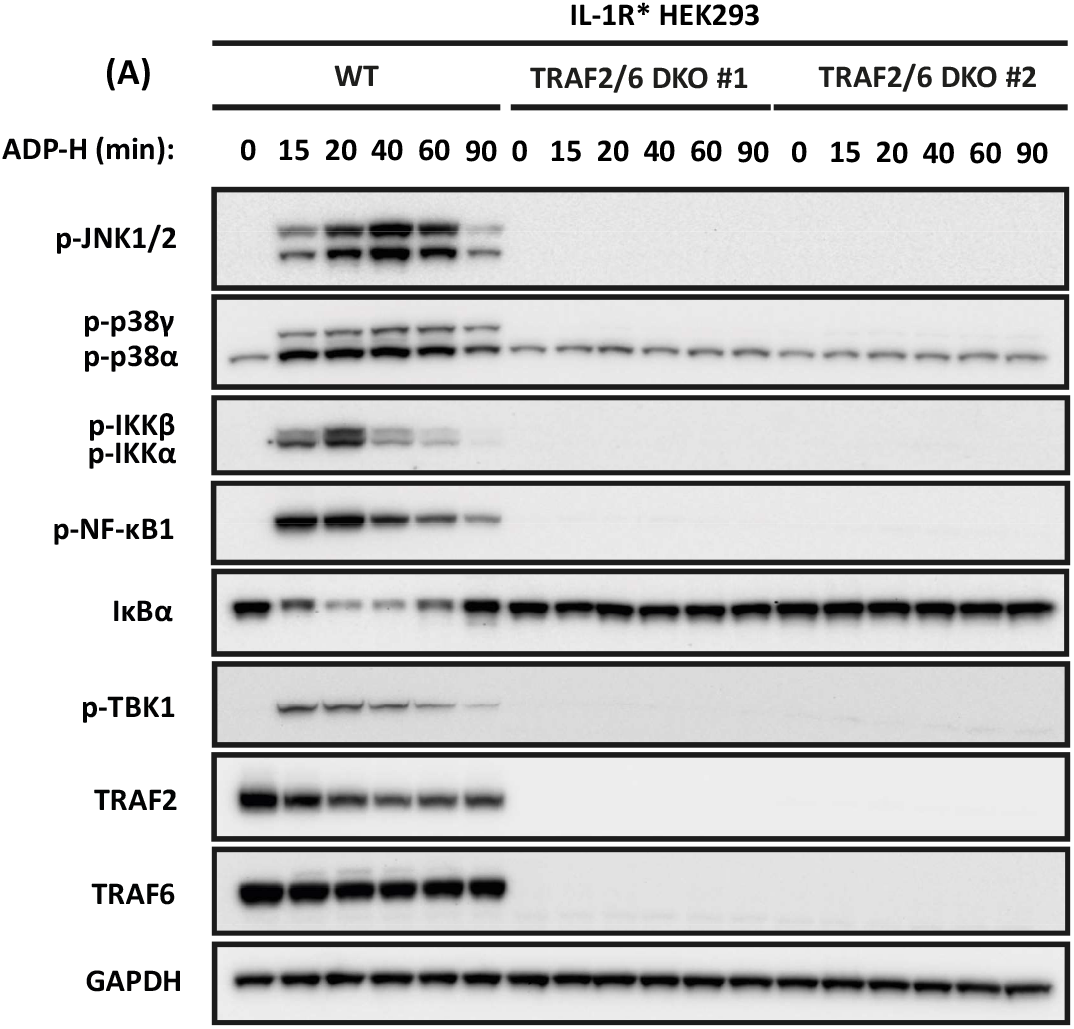
ADP-heptose signalling is disrupted in cells lacking TRAF2 and TRAF6. Parental and two different clones of TRAF2/TRAF6 double KO IL-1R* HEK293 cells were stimulated with ADP-H for the times indicated. The cell extracts were subjected SDS-PAGE and immunoblotting with the antibodies indicated. ADP-H did not increase the basal level of TBK1 phosphorylation significantly in the TRAF2/TRAF6 double knockout cells.

## References

Beinke, S., Robinson, M.J., Hugunin, M. and Ley, S.C. (2004) Lipopolysaccharide activation of the TPL-2/MEK/extracellular signal-regulated kinase mitogen-activated protein kinase cascade is regulated by IkappaB kinase-induced proteolysis of NF-kappaB1 p105, Mol Cell Biol., 24, 9658–9667.

Boulard, O., Kirchberger, S., Royston, D.J., Maloy, K.J. and Powrie, F.M. (2012) Identification of a genetic locus controlling bacteria-driven colitis and associated cancer through effects on innate inflammation, J Exp Med., 209, 1309–1324.

Campbell, D.G. and Morrice, N.A. (2002) Identification of protein phosphorylation sites by a combination of mass spectrometry and solid phase Edman sequencing, J Biomol Tech., 13, 119–130.

Clark, K., Peggie, M., Plater, L., Sorcek, R.J., Young, E.R.R., Madwed, J.B., Hough, J., McIver, E.G. and Cohen, P. (2011) Novel cross-talk within the IKK family controls innate immunity, Biochem J., 434, 93–104.

Cohen, P. and Strickson, S. (2017) The role of hybrid ubiquitin chains in the MyD88 and other innate immune signalling pathways, Cell Death & Differentiation, 24, 1153–1159.

Delhase, M., Makio, H., Chen, Y. and Karin, M. (1999) Positive and negative regulation of IκB kinase activity through IKKβ subunit phosphorylation, Science, 284, 309–313.

Duckett, C.S. (2005) IAP proteins: sticking it to Smac, Biochem J., 385, 1–2.

Ea, C.K., Sun, L., Inoue, J.I. and Chen, Z.J. (2004) TIFA activates IκB kinase (IKK) by promoting oligomerization and ubiquitination of TRAF6, Natl Acad Sci U S A., 101, 15318– 15323.

Emmerich, C.H. and Cohen, P. (2015) Optimising methods for the preservation, capture and identification of ubiquitin chains and ubiquitylated proteins by immunoblotting, Biochem Biophys Res Commun., 466, 1–14.

Emmerich, C.H., Ordureau, A., Strickson, S., Arthur, J.S.C., Pedrioli, P.G.A.A., Komander, D. and Cohen, P. (2013) Activation of the canonical IKK complex by K63/M1-linked hybrid ubiquitin chains, Proc Natl Acad Sci U S A., 110, 15247–15252.

Fang, C.Y., Chen, H.Y., Wang, M., Chen, P.L., Chang, C.F., Chen, L.S., Shen, C.H., Ou, W.C., Tsai, M.D., Hsu, P.H. and Chang, D. (2010) Global analysis of modifications of the human BK virus structural proteins by LC-MS/MS, Virology, 402, 164–176.

Gall, A., Gaudet, R.G., Gray-Owen, S.D. and Salama, N.R. (2017) TIFA signaling in gastric epithelial cells initiates the cag type 4 secretion system-dependent innate immune response to helicobacter pylori infection, mBio, 8, e01168–17.

Gaudet, R.G., Sintsova, A., Buckwalter, C.M., Leung, N., Cochrane, A., Li, J., Cox, A.D., Moffat, J. and Gray-Owen, S.D. (2015) Cytosolic detection of the bacterial metabolite HBP activates TIFA-dependent innate immunity, Science, 348, 1251–1255.

Goh, E.T.H., Arthur, J.S.C., Cheung, P.C.F., Akira, S., Toth, R. and Cohen, P. (2012) Identification of the protein kinases that activate the E3 ubiquitin ligase Pellino 1 in the innate immune system, Biochem J., 441, 339–346.

Huang, C.-C.F., Weng, J.-H., Wei, T.-Y.W., Wu, P.-Y.G., Hsu, P.-H., Chen, Y.-H., Wang, S.-C., Qin, D., Hung, C.-C., Chen, S.-T., Wang, A.H.-J., Shyy, J.Y.-J. and Tsai, M.-D. (2012) Intermolecular binding between TIFA-FHA and TIFA-pT mediates tumor necrosis factor alpha stimulation and NF-κB activation, Mol Cell Biol., 32, 3800–3800.

Kanamori, M., Suzuki, H., Saito, R., Muramatsu, M. and Hayashizaki, Y. (2002) T2BP, a novel TRAF2 binding protein, can activate NF-κB and AP-1 without TNF stimulation, Biochem Biophys Res Commun., 290, 1108–1113.

Kanayama, A., Seth, R.B., Sun, L., Ea, C.K., Hong, M., Shaito, A., Chiu, Y.H., Deng, L. and Chen, Z.J. (2004) TAB2 and TAB3 activate the NF-κB pathway through binding to polyubiquitin chains, Mol Cell., 15, 535–548.

Kirisako, T., Kamei, K., Murata, S., Kato, M., Fukumoto, H., Kanie, M., Sano, S., Tokunaga, F., Tanaka, K. and Iwai, K. (2006) A ubiquitin ligase complex assembles linear polyubiquitin chains, EMBO J., 25, 4877–4887.

Kishore, N., Khai Huynh, Q., Mathialagan, S., Hall, T., Rouw, S., Creely, D., Lange, G., Caroll, J., Reitz, B., Donnelly, A., Boddupalli, H., Combs, R.G., Kretzmer, K. and Tripp, C.S. (2002) IKK-i and TBK-1 are enzymatically distinct from the homologous enzyme IKK-2: Comparative analysis of recombinant human IKK-i, TBK-1, and IKK-2, J Biol Chem., 277, 13840–13847.

Kulathu, Y., Akutsu, M., Bremm, A., Hofmann, K. and Komander, D. (2009) Two-sided ubiquitin binding explains specificity of the TAB2 NZF domain, Nat Struct Mol Biol., 16, 1328–1330.

Li, W., Li, B., Giacalone, N.J., Torossian, A., Sun, Y., Niu, K., Lin-Tsai, O. and Lu, B. (2011) BV6, an IAP antagonist, activates apoptosis and enhances radiosensitization of non-small cell lung carcinoma in vitro, J Thorac Oncol., 6, 1801.

Lo, Y.C., Lin, S.C., Rospigliosi, C.C., Conze, D.B., Wu, C.J., Ashwell, J.D., Eliezer, D. and Wu, H. (2009) Structural basis for recognition of diubiquitins by NEMO, Mol Cell., 33, 602– 615.

Mahajan, A., Yuan, C., Lee, H., Chen, E.S.W., Wu, P.Y. and Tsai, M.D. (2008) Structure and function of the phosphothreonine-specific FHA domain, Sci Signal., 1, re12.

Maubach, G., Lim, M.C.C., Sokolova, O., Backert, S., Meyer, T.F. and Naumann, M. (2021) TIFA has dual functions in Helicobacter pylori-induced classical and alternative NF-κB pathways, EMBO Rep., 22, e52878.

Medzhitov, R. (2007) Recognition of microorganisms and activation of the immune response, Nature, 449, 819–826.

Middleton, A.J., Budhidarmo, R., Das, A., Zhu, J., Foglizzo, M., Mace, P.D. and Day, C.L. (2017) The activity of TRAF RING homo- and heterodimers is regulated by zinc finger 1, Nat Commun., 8, 1–10.

Milivojevic, M., Dangeard, A.S., Kasper, C.A., Tschon, T., Emmenlauer, M., Pique, C., Schnupf, P., Guignot, J. and Arrieumerlou, C. (2017) ALPK1 controls TIFA/TRAF6-dependent innate immunity against heptose-1,7-bisphosphate of gram-negative bacteria, PLoS Pathog., 13, e1006224.

Ninomiya-Tsuji, J., Kajino, T., Ono, K., Ohtomo, T., Matsumoto, M., Shiina, M., Mihara, M., Tsuchiya, M. and Matsumoto, K. (2003) A resorcylic acid lactone, 5Z-7-oxozeaenol, prevents inflammation by inhibiting the catalytic activity of TAK1 MAPK kinase kinase, J Biol Chem., 278, 18485–18490.

Ordureau, A., Smith, H., Windheim, M., Peggie, M., Carrick, E., Morrice, N. and Cohen, P. (2008) The IRAK-catalysed activation of the E3 ligase function of Pellino isoforms induces the Lys63-linked polyubiquitination of IRAK1, Biochem J., 409, 43–52.

Palkowitsch, L., Leidner, J., Ghosh, S. and Marienfeld, R.B. (2008) Phosphorylation of serine 68 in the IkappaB kinase (IKK)-binding domain of NEMO interferes with the structure of the IKK complex and tumor necrosis factor-alpha-induced NF-kappaB activity, J Biol Chem., 283, 76–86.

Pauls, E., Nanda, S.K., Smith, H., Toth, R., Arthur, J.S.C. and Cohen, P. (2013) Two phases of inflammatory mediator production defined by the study of IRAK2 and IRAK1 knock-in mice, J Immunol., 191, 2717–2730.

Perez-Riverol, Y., Bai, J., Bandla, C., García-Seisdedos, D., Hewapathirana, S., Kamatchinathan, S., Kundu, D.J., Prakash, A., Frericks-Zipper, A., Eisenacher, M., Walzer, M., Wang, S., Brazma, A. and Vizcaíno, J.A. (2022) The PRIDE database resources in 2022: a hub for mass spectrometry-based proteomics evidences, Nucleic Acids Res., 50, D543–D552.

Qin, J.Y., Zhang, L., Clift, K.L., Hulur, I., Xiang, A.P., Ren, B.Z. and Lahn, B.T. (2010) Systematic comparison of constitutive promoters and the Doxycycline-inducible promoter, PLoS One., 5, e10611.

Rahighi, S., Ikeda, F., Kawasaki, M., Akutsu, M., Suzuki, N., Kato, R., Kensche, T., Uejima, T., Bloor, S., Komander, D., Randow, F., Wakatsuki, S. and Dikic, I. (2009) Specific recognition of linear ubiquitin chains by NEMO is important for NF-kappaB activation, Cell, 136, 1098–1109.

Ryzhakov, G., West, N.R., Franchini, F., Clare, S., Ilott, N.E., Sansom, S.N., Bullers, S.J., Pearson, C., Costain, A., Vaughan-Jackson, A., Goettel, J.A., Ermann, J., Horwitz, B.H., Buti, L., Lu, X., Mukhopadhyay, S., Snapper, S.B. and Powrie, F. (2018) Alpha kinase 1 controls intestinal inflammation by suppressing the IL-12/Th1 axis, Nat Commun., 9, 1–13.

Schaeffer, V., Akutsu, M., Olma, M.H., Gomes, L.C., Kawasaki, M. and Dikic, I. (2014) Binding of OTULIN to the PUB domain of HOIP controls NF-κB signaling, Mol Cell., 54, 349–361.

Schäffer, C. and Messner, P. (2004) Surface-layer glycoproteins: an example for the diversity of bacterial glycosylation with promising impacts on nanobiotechnology, Glycobiology, 14, 31R–42R.

Shevchenko, A., Tomas, H., Havliš, J., Olsen, J. V. and Mann, M. (2007) In-gel digestion for mass spectrometric characterization of proteins and proteomes, Nat Protoc., 1, 2856–2860.

Smith, H., Peggie, M., Campbell, D.G., Vandermoere, F., Carrick, E. and Cohen, P. (2009) Identification of the phosphorylation sites on the E3 ubiquitin ligase Pellino that are critical for activation by IRAK1 and IRAK4, Proc Natl Acad Sci U S A., 106, 4584–4590.

Stein, S.C., Faber, E., Bats, S.H., Murillo, T., Speidel, Y., Coombs, N. and Josenhans, C. (2017) Helicobacter pylori modulates host cell responses by CagT4SS-dependent translocation of an intermediate metabolite of LPS inner core heptose biosynthesis, PLoS Pathog., 13, e1006514.

Strickson, S., Emmerich, C.H., Goh, E.T.H., Zhang, J., Kelsall, I.R., MacArtney, T., Hastie, C.J., Knebel, A., Peggie, M., Marchesi, F., Arthur, J.S.C. and Cohen, P. (2017) Roles of the TRAF6 and Pellino E3 ligases in MyD88 and RANKL signaling, Proc Natl Acad Sci U S A., 114, E3481–E3489.

Sun, S.C. (2017) The non-canonical NF-κB pathway in immunity and inflammation, Nat Rev Immunol., 17, 545–558.

Tada, K., Okazaki, T., Sakon, S., Kobarai, T., Kurosawa, K., Yamaoka, S., Hashimoto, H., Mak, T.W., Yagita, H., Okumura, K., Yeh, W.C. and Nakano, H. (2001) Critical roles of TRAF2 and TRAF5 in tumor necrosis factor-induced NF-kappa B activation and protection from cell death, J Biol Chem., 276, 36530–36534.

Takatsuna, H., Kato, H., Gohda, J., Akiyama, T., Moriya, A., Okamoto, Y., Yamagata, Y., Otsuka, M., Umezawa, K., Semba, K. and Inoue, J. (2003) Identification of TIFA as an adapter protein that links tumor necrosis factor receptor-associated factor 6 (TRAF6) to Interleukin-1 (IL-1) receptor-associated kinase-1 (IRAK-1) in IL-1 receptor signaling, J Biol Chem., 278, 12144–12150.

Tan, L. et al. (2015) Discovery of type II inhibitors of TGFβ-activated kinase 1 (TAK1) and mitogen-activated protein kinase kinase kinase kinase 2 (MAP4K2), J. Med. Chem., 58, 183– 196.

Tang, W., Guo, Z., Cao, Z., Wang, M., Li, P., Meng, X., Zhao, X., Xie, Z., Wang, W., Zhou, A., Lou, C. and Chen, Y. (2018) D-Sedoheptulose-7-phosphate is a common precursor for the heptoses of septacidin and hygromycin B, Proc Natl Acad Sci U S A., 115, 2818–2823.

Tokunaga, F., Sakata, S.I., Saeki, Y., Satomi, Y., Kirisako, T., Kamei, K., Nakagawa, T., Kato, M., Murata, S., Yamaoka, S., Yamamoto, M., Akira, S., Takao, T., Tanaka, K. and Iwai, K. (2009) Involvement of linear polyubiquitylation of NEMO in NF-κB activation, Nature Cell Biology 2009 11:2, 11, 123–132.

Walsh, M.C., Kim, G.K., Maurizio, P.L., Molnar, E.E. and Choi, Y. (2008) TRAF6 autoubiquitination-independent activation of the NFκB and MAPK pathways in response to IL-1 and RANKL, PLoS One., 3, e4064.

Wang, C., Deng, L., Hong, M., Akkaraju, G.R., Inoue, J.I. and Chen, Z.J. (2001) TAK1 is a ubiquitin-dependent kinase of MKK and IKK, Nature, 412, 346–351.

Wang, S. (2010) Design of small-molecule Smac mimetics as IAP antagonists, Curr Top Microbiol Immunol., 348, 89–113.

Waterfield, M., Jin, W., Reiley, W., Zhang, M. and Sun, S.-C. (2004) IκB kinase is an essential component of the Tpl2 signaling pathway, Mol Cell Biol., 24, 6040–6048.

Yamamoto, Y., Yin, M.-J. and Gaynor, R.B. (2000) IkappaB kinase alpha (IKKalpha) regulation of IKKbeta kinase activity, Mol Cell Biol., 20, 3655–3666.

Ye, H., Arron, J.R., Lamothe, B., Cirilli, M., Kobayashi, T., Shevde, N.K., Segal, D., Dzivenu, O.K., Vologodskaia, M., Yim, M., Du, K., Singh, S., Pike, J.W., Darnay, B.G., Choi, Y. and Wu, H. (2002) Distinct molecular mechanism for initiating TRAF6 signalling, Nature, 418, 443–447.

Ye, H., Park, Y.C., Kreishman, M., Kieff, E. and Wu, H. (1999) The structural basis for the recognition of diverse receptor sequences by TRAF2, Mol Cell., 4, 321–330.

Yin, Q., Lin, S.-C., Lamothe, B., Lu, M., Lo, Y.-C., Hura, G., Zheng, L., Rich, R.L., Campos, A.D., Myszka, D.G., Lenardo, M.J., Darnay, B.G. and Wu, H. (2009) E2 interaction and dimerization in the crystal structure of TRAF6, Nat Struct Mol Biol., 16, 658–666.

Zamyatina, A., Gronow, S., Puchberger, M., Graziani, A., Hofinger, A. and Kosma, P. (2003) Efficient chemical synthesis of both anomers of ADP l-glycero- and d-glycero-d-manno-heptopyranose, Carbohydr Res., 338, 2571–2589.

Zhang, J., Clark, K., Lawrence, T., Peggie, M.W., Cohen, P., Zhang, J., Cohen, P. and Peggie, M.W. (2014) An unexpected twist to the activation of IKKβ: TAK1 primes IKKβ for activation by autophosphorylation, Biochem J., 461, 531–537.

Zhang, J., Macartney, T., Peggie, M. and Cohen, P. (2017) Interleukin-1 and TRAF6-dependent activation of TAK1 in the absence of TAB2 and TAB3, Biochem J., 474, 2235– 2248.

Zhou, P., She, Y., Dong, N., Li, P., He, H., Borio, A., Wu, Q., Lu, S., Ding, X., Cao, Y., Xu, Y., Gao, W., Dong, M., Ding, J., Wang, D.C., Zamyatina, A. and Shao, F. (2018) Alpha-kinase 1 is a cytosolic innate immune receptor for bacterial ADP-heptose, Nature, 561, 122–126.

Zimmermann, S. et al. (2017) ALPK1- and TIFA-dependent innate immune response triggered by the Helicobacter pylori Type IV secretion system, Cell Rep., 20, 2384–2395.

